# Immature myeloid cells are indispensable for intestinal regeneration post irradiation injury

**DOI:** 10.1101/2023.02.28.530500

**Authors:** Zhengyu Jiang, Quin T. Waterbury, Na Fu, Woosook Kim, Ermanno Malagola, Chandan Guha, Carrie J. Shawber, Kelley S. Yan, Timothy C. Wang

**Affiliations:** Division of Digestive and Liver Diseases Medicine and Irving Cancer Research Center, Department of Medicine, Columbia University Medical Center, New York, NY 10032, USA; Janssen Research and Development, Johnson & Johnson, PA 19044, USA; Institute of Human Nutrition, Columbia University Medical Center, New York, NY 10032, USA; Department of Traditional and Western Medical Hepatology, Third Hospital of Hebei Medical University, Shijiazhuang, Hebei, China; Department of Radiation Oncology, Montefiore Medical Center, Bronx, NY 10467, USA; Department of Pathology, Albert Einstein College of Medicine, Bronx, NY 10467, USA; Deparments of Obstetrics and Gynecology and Surgery, Columbia University Irving Medical Center, New York, NY 10032, USA; Department of Genetics and Development, Columbia University Medical Center, New York, NY, 10032, USA

**Keywords:** Immature myeloid cell, lymphatic, endothelial cell, epithelium, PGE2, RSPO3, microbiome, intestine, regeneration

## Abstract

The intestinal epithelium functions both in nutrient absorption and as a barrier, separating the luminal contents from a network of vascular, fibroblastic, and immune cells underneath. Following injury to the intestine, multiple different cell populations cooperate to drive regeneration of the mucosa. Immature myeloid cells (IMCs), marked by histidine decarboxylase (*Hdc*), participate in regeneration of multiple organs such as the colon and central nervous system. Here, we found that IMCs infiltrate the injured intestine and promote epithelial regeneration and modulate LEC activity. IMCs produce prostaglandin E2 (PGE2), which promotes LEC lymphangiogenesis and upregulation of pro-regenerative factors including RSPO3. Moreover, we found that IMC recruitment into the intestine is driven by invading microbial signals. Accordingly, antibiotic eradication of the intestinal microbiome prior to WB-IR inhibits IMC recruitment, and consequently, intestinal recovery. We propose that IMCs play a critical role in intestinal repair and implicate gut microbes as mediators of intestinal regeneration.

## INTRODUCTION

Epithelial tissues such as the skin or intestine undergo constant stress and injury and must continuously regenerate to maintain proper barrier function. Compromise of the intestinal barrier leads to penetration of numerous microbial products from commensal bacteria, such as the endotoxin class lipopolysaccharides (LPS), which affects tissues locally and systemically. LPS exposure is generally considered harmful, so regeneration of the epithelium is tightly regulated to limit endotoxin exposure and maintain proper healing after injury without excessive growth (Blanpain and Fuchs, 2014). In the intestine, regeneration of the intestinal epithelium relies on complex crosstalk between highly active intestinal stem cells (ISCs) residing near the crypt base, and surrounding niche cells that include endothelial cells, fibroblasts, and immune cells (Antanaviciute et al., 2022; McCarthy et al., 2020a; Saha et al., 2016). These niche cells produce several signals, including R-spondins, Wns, and Egf family ligands, that are essential for ISC maintenance and epithelial regeneration (Abud et al., 2005; Jarde et al., 2020; McCarthy et al., 2020b; Yan et al., 2017).

Recently, lymphatic endothelial cells (LECs) have been shown to be a major source of R-spondin 3 (RSPO3) (Niec et al., 2022; Ogasawara et al., 2018). RSPO3 is the major ligand for the Lgr4/5 receptors on ISCs and progenitor cells and is necessary for proper regeneration after injury (Harnack et al., 2019). LECs exist in a network in the intestine that consists of a central lacteal located in the center of the villus, which connects to pre-collectors in the submucosa (Choe et al., 2015). Pre-collector LECs surround the crypt base and associate with ISCs to regulate their activity (Niec et al., 2022). This LEC population expands in the intestine after whole-body irradiation (WB-IR), and conditional knockout of the *Rspo3* gene in LECs hinders intestinal regeneration after WB-IR or 5FU chemotoxic injury, highlighting the importance of LEC-derived RSPO3 in this setting (Goto et al., 2022; Palikuqi et al., 2022). Lymphatics also expand in the bone after WB-IR and participate in regeneration (Biswas et al., 2023). While the expansion of LECs and increased RSPO3 production is necessary for intestinal regeneration, it is unclear how LECs are activated and RSPO3 secretion is induced in response to intestinal injury.

Myeloid cells have been shown to play essential roles in intestinal homeostasis and regeneration (Bain and Mowat, 2014; De Schepper et al., 2018; Shaw et al., 2018). During injury, Ly6C^+^ monocytes rapidly accumulate in the intestine and integrate signals coming from gut microbes to the injured epithelium through toll-like receptor (TLR) signaling (Kim et al., 2018; Smith et al., 2011). Infiltrating monocytes are a vital source of Wnts that promote survival of Lgr5^+^ cells during injury and epithelial repair (Saha et al., 2016). Myeloid-derived Wnt signaling also plays an important role in regulating LEC proliferation in the dermis (Muley et al., 2017). Myeloid cells can also regulate lymphatic cell angiogenesis through the production of VEGF-A and VEGF-C (Cursiefen et al., 2004; Machnik et al., 2009). In addition to Wnt and Vegf signaling, myeloid cells in the intestine also express *Ptgs2* (the gene encoding the COX-2 enzyme) and are known to be a source of prostaglandin E2 (PGE2) (Ishikawa et al., 2011). PGE2 is a bioactive lipid that mediates inflammatory signals through several different types of receptors, including the E-type prostanoid receptors (EP) (Hull et al., 2004). PGE2 has been known to protect mice against IR-injury induced death since the 1980s (Walden et al., 1987). Specifically, pretreating mice with PGE2 protects epithelial cells from IR-induced apoptosis by inhibiting Bax signaling (Tessner et al., 2004). Moreover, PGE2 is also able to promote intestinal epithelium repair by expanding the pool of regenerative progenitor cells through EP4 receptor signaling (Ishikawa et al., 2011; Miyoshi et al., 2017; Roulis et al., 2020). Notably, in breast tissue, PGE2 is a known activator of LEC lymphangiogenesis through the EP4 receptor, but its role in regulating intestinal LECs is unknown (Nandi et al., 2017).

Histidine decarboxylase (HDC) generates endogenous histamine from dietary L-histidine and marks a lineage of hematopoietic stem cells (HSCs) and immature myeloid cells (Chen et al., 2017). Hdc^+^ immature myeloid cells (IMCs) are primarily granulocytic, marked by CD11b^+^ Ly6G^+^. During homeostasis, these cells comprise ≥80% of IMCs in the bone marrow and are rarely found in the circulation or peripheral tissues (Fu et al., 2021). In addition to maintaining the HSC niche through histamine production, this IMC population is rapidly mobilized from the bone marrow to damaged tissues in response to inflammatory conditions such as acute colitis or LPS endotoxemia, where they contribute to growth and regeneration (Chen et al., 2017; Fu et al., 2021).

Here, we studied the role of Hdc^+^ IMCs in intestinal regeneration after IR injury. We found that following damage to the intestinal barrier, LECs are exposed to microbial-derived LPS, which triggers their upregulation of *Cxcl1*. This in turn recruits IMCs near intestinal LECs in a CXCR2-dependent manner. Within the intestine, IMCs support epithelial regeneration by secreting PGE2, which activates the LECs and promotes their upregulation of pro-regenerative factors, including *Rspo3*, *Ccl21a*, and *Wnt2a*. Using transgenic mouse models, we show that IMCs are essential for mediating the response to microbial signals and orchestrating LECs’ role in intestinal regeneration.

## RESULTS

### Hdc+ IMCs are recruited to the damaged intestine following irradiation injury

We have previously shown that Hdc^+^ IMCs are mobilized from the bone marrow to the colon during DSS-induced colitis (Fu et al., 2021). To test if Hdc^+^ IMCs are involved in intestinal regeneration, we employed a model of 12 Gy whole-body irradiation (WB-IR), which induces acute damage to the intestinal epithelial stem and progenitor zone cells (Booth et al., 2012). To track the movement of IMCs, we took advantage of our previously generated Hdc^GFP^ mice, which label the majority of CD11b^+^ Gr-1^+^ IMCs (Yang et al., 2011). Immunofluorescent staining and flow cytometry analyses revealed that Hdc^GFP+^ cells are virtually absent from the intestine of unirradiated mice but infiltrate the tissue one day after 12 Gy WB-IR exposure, reaching maximum abundance at 3 days post WB-IR (**Figures 1A – D, Supplementary Figures S1A – D**). Within 10 days post-IR, when regeneration is mostly finished (Liang et al., 2020), Hdc^GFP+^ cells were lost from the intestine. Importantly, immunophenotypic analysis confirmed these Hdc^GFP+^ cells in the intestine are CD45^+^ CD11b^+^ Ly6G^intermediate^ granulocytic IMCs, as opposed to Ly6C+ monocytes (**Supplementary Figure S1E**).

**Figure 1:**
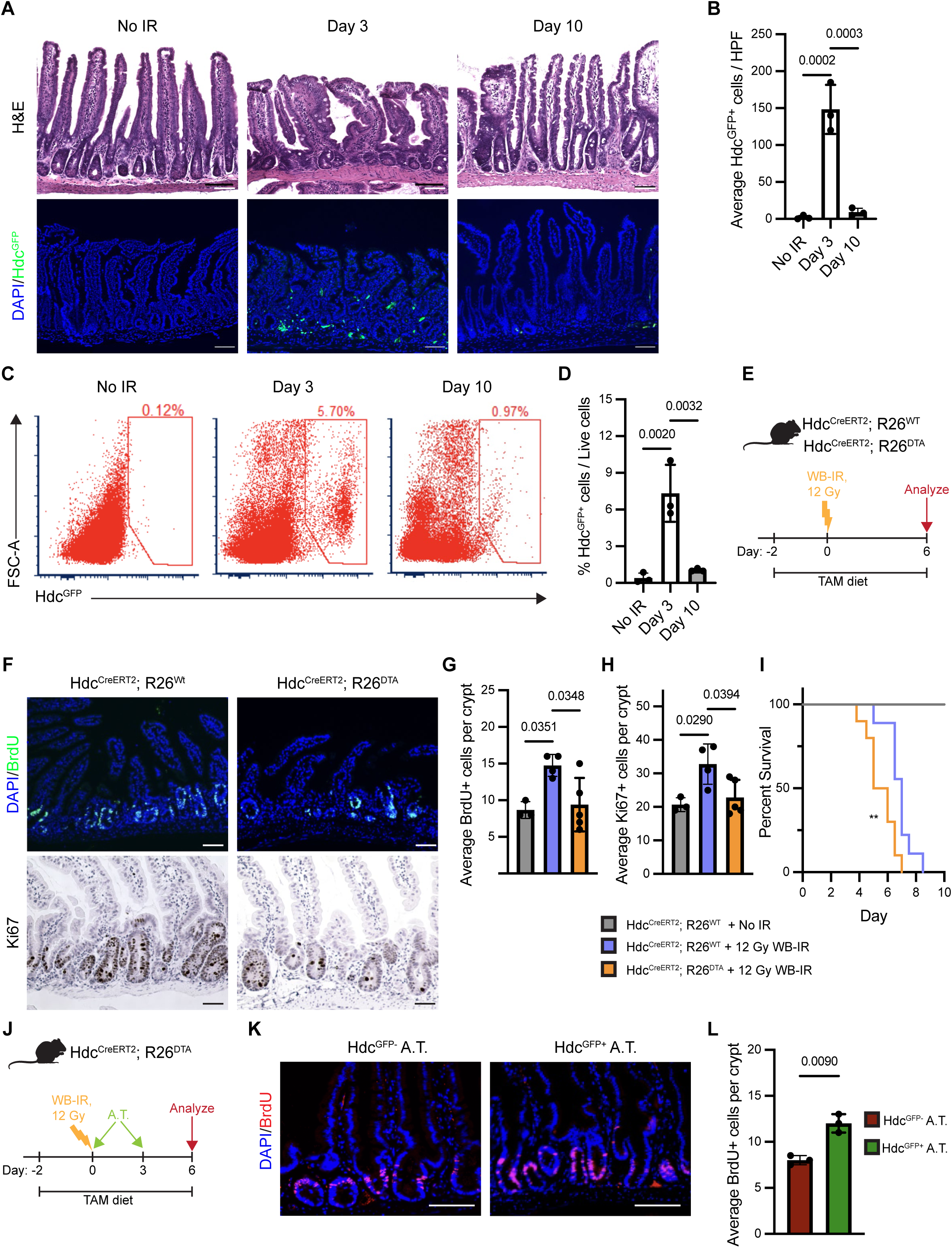
Hdc+ IMCs are indispensable for intestinal regeneration after IR. **A.** Representative 100x images of H&E staining (top) and GFP visualization (bottom) from the proximal jejunum of non-irradiated Hdc^GFP^ mice or mice 3 and 10 days after 12 Gy WB-IR. **B.** Quantification of GFP+ cells of images in (**A**). (n=3), HPF= high powered field. **C.** Representative FACS plots of live (DAPI^-^) intestinal cells from non-irradiated mice or mice 3 and 10 days after 12 Gy WB-IR. **D.** Bar plot showing quantification of (**C**). (n=3). **E.** Experimental scheme of Hdc+ cell ablation experiment. TAM diet = tamoxifen diet. **F.** Representative 100x images of BrdU (top) and Ki67 (bottom) staining from the proximal jejunum of Hdc^CreERT2^; R26^WT^ (control) or Hdc^CreERT2^; R26^DTA^ mice after 12 Gy WB-IR. **G.** Bar plot showing quantification of BrdU staining from (**E**) (n=3-5). **H.** Bar plot showing quantification of Ki67 staining from (**E**). (n=3-5). **I.** Survival curve of Hdc^CreERT2^ (control) or Hdc^CreERT2^; R26^DTA^ mice receiving a tamoxifen diet with or without 12 Gy WB-IR. Hdc^CreERT2^; R26^DTA^ no IR median: >10 days. Hdc^CreERT2^ with IR median: 7 days. Hdc^CreERT2^; R26^DTA^ with IR median: 5.75 days. (n=5, 9 or 10), **: p<0.005. **J.** Experimental scheme of adoptive transfer study. A.T.= adoptive transfer. **K.** Representative 100x images of BrdU immunostaining from the proximal jejunum of Hdc^CreERT2^; R26^DTA^ mice that received adoptive transfer of Hdc^GFP-^ bone marrow cells (GFP^-^ AT) or Hdc^GFP+^ bone marrow cells (GFP^+^ AT) from healthy Hdc^GFP^ mice twice after 12 Gy WB-IR. **L.** Bar plot showing quantification of BrdU staining from (**J**). (n=3). Scale bars = 50 um Bar graph data are mean ± SEM. Statistical analysis was performed using an Ordinary one-way ANOVA with multiple comparisons to each group (**B, D, G, H**), Kaplan-Meier simple survival analysis (**I**) or a two-sided Student’s t test (**L**).

### Hdc+ IMCs are indispensable for intestinal regeneration after IR

Since the gut-homing kinetics of Hdc^+^ IMCs following IR aligns with the dynamics of intestinal regeneration, we hypothesized that IMCs may be important for intestinal regeneration after IR. To test this hypothesis, we selectively ablated Hdc^+^ cells *in vivo* using Hdc^CreERT2^; R26^DTA^ mice (Chen et al., 2017). Following Cre induction via a tamoxifen-containing diet, Hdc^CreERT2^; R26^DTA^ mice showed an up to 10-fold reduction in IMCs infiltrating the intestine 3 days post-IR, relative to controls (Hdc^CreERT2^; R26^WT^), confirming the cell depletion in this model (**Figure 1E**) (**Supplementary Figure S1F – H**). Ablation of Hdc+ cells in the setting of IR injury reduced intestinal regeneration, as evidenced by fewer BrdU+ and Ki67+ cells per crypt in Hdc^CreERT2^; R26-DTA mice relative to controls (**Figures 1F – 1H**). This impairment of intestinal regeneration was associated with decreased survival (**Figure 1I**). Importantly, adoptive transfer of Hdc^GFP+^, but not Hdc^GFP-^, bone marrow cells from non-irradiated Hdc^GFP^ mice into irradiated Hdc^CreERT^; R26^DTA^ mice partially rescued intestinal regeneration (**Figures 1J – L**). Together, these results suggest that Hdc+ IMCs are indispensable for proper intestinal regeneration after IR injury.

### Hdc+ IMCs promote epithelial regeneration via production of PGE2

Myeloid cells in the intestine are known to be sources of PGE2 (Crittenden et al., 2021; Ishikawa et al., 2011), which has been shown to promote intestinal epithelial regeneration (Miyoshi et al., 2017; Roulis et al., 2020). We found that Hdc^+^ IMCs express *Ptgs2*, encoding COX-2, a key enzyme responsible for PGE2 production. Interestingly, intestinally recruited Hdc^+^ IMCs express significantly higher levels of *Ptgs2* compared to those found in the bone marrow of healthy mice (**Figure 2A**). To determine if IMC-derived PGE2 is important for intestinal regeneration post IR exposure, we selectively abrogated *Ptgs2* expression in Hdc^+^ cells using Hdc^Cre^; *Ptgs2*^fl/fl^ mice (Ishikawa and Herschman, 2006). Following 12 Gy WB-IR challenge, Hdc^Cre^; *Ptgs2*^fl/fl^ mice showed significantly fewer proliferating BrdU+ cells per crypt and reduced villus length compared to Hdc^Cre^ controls, similar to what we observed in the setting of Hdc^+^ cell ablation (**Figures 2B – E**).

**Figure 2:**
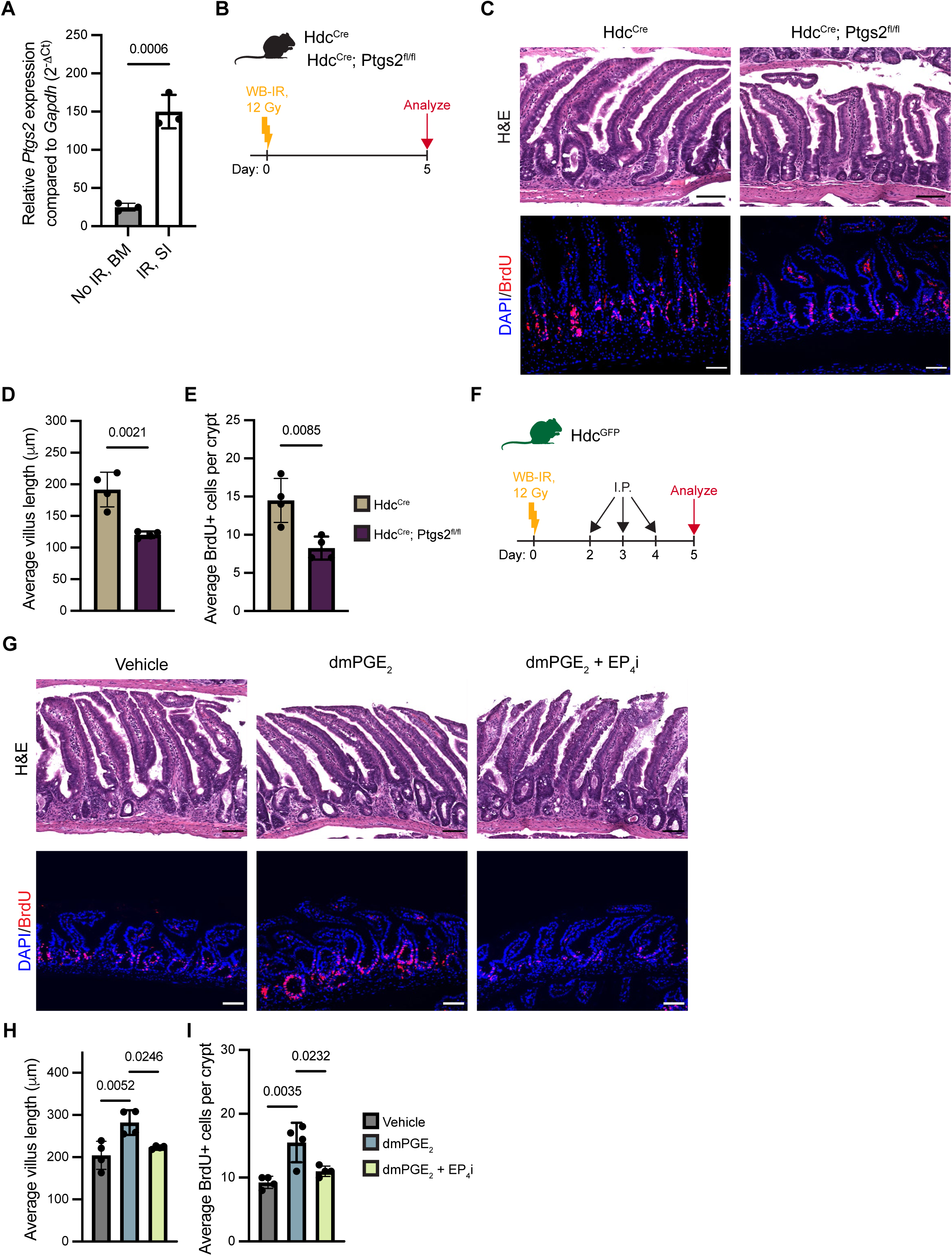
Hdc+ IMCs produce PGE2 that promotes intestinal regeneration. **A)** qPCR for *Ptgs2* in Hdc^+^ IMCs isolated from the bone marrow of healthy Hdc^GFP^ mice (no IR, BM) and intestine of mice 3 days after receiving 12 Gy WB-IR (IR, SI). (n=3). **B.** Experimental scheme of conditional *Ptgs2* knockout irradiation experiment. **C.** Representative 100x images of H&E staining (top) and BrdU immunostaining (bottom) from the proximal jejunum of Hdc^Cre^ (control) or Hdc^Cre^; Ptgs2^fl/fl^ mice after receiving 12 Gy WB-IR. **D.** Bar plot showing quantification of the average villus length from (**B**). (n=4). **E.** Bar plot showing quantification of BrdU staining from (**B**). (n=4). **F.** Experimental scheme of dmPGE2 treatment after irradiation experiment. I.P. = intraperitoneal injection of vehicle, dmPGE2, or dmPGE2 with EP4i pretreatment. **G.** Representative images of H&E staining (top) and BrdU immunostaining (bottom) from the proximal jejunum of Hdc^GFP^ mice treated with vehicle, dmPGE2, or dmPGE2 + EP4i pretreatment after 12 Gy WB-IR. **H.** Bar plot showing quantification of the average villus length from (**F**) (n=4). **I.** Bar plot showing quantification of BrdU staining from (**F**). (n=4). Scale bars = 50 uM Bar graph data are mean ± SEM. Statistical analysis was performed using a two-sided Student’s t test (**A, D, E**) or an Ordinary one-way ANOVA with multiple comparisons to each group (**H, I**).

To further investigate the role of PGE2 in epithelial cell proliferation, we treated intestinal organoids with the stable PGE2 analog 16,16 dimethylprostaglandin E2 (dmPGE2) *in vitro*. Organoids treated with dmPGE2 showed increased diameter but not number, suggesting PGE2 may act as a growth factor for epithelial cells, consistent with previous literature (Ishikawa et al., 2011; Miyoshi et al., 2017). The effects of dmPGE2 on organoids was reversed by adding a selective inhibitor for the prostaglandin receptor EP4 (EP4i) (**Supplementary Figures S2A – C**). Given the importance of Hdc^+^ IMC-derived PGE2 in intestinal epithelial regeneration following IR, we tested the effects of dmPGE2 *in vivo* on mice challenged with 12 Gy WB-IR. To distinguish from the reported function of dmPGE2 in protecting epithelial cells against IR-induced apoptosis, we began treatment two days after IR, when apoptosis of epithelial cells has already occurred (Booth et al., 2012; Tessner et al., 2004). Five days after irradiation, dmPGE2-treated mice had increased villus length and BrdU+ cells per crypt compared to vehicle treated mice, but these effects could be abrogated by pretreatment with EP4i (**Figures 2F – I**). Intriguingly, unirradiated mice that received dmPGE2 also showed more BrdU+ cells per crypt, but no difference in villus length, suggesting that the pro-mitogenic activity of PGE2 is independent of tissue damage (**Supplementary Figures S2D – H**).

Altogether, these data indicate that Hdc+ IMCs promote intestinal regeneration after IR, at least in part, by secreting PGE2, which activates epithelial proliferation through EP4 signaling.

### Hdc+ IMCs activate LECs via a PGE2/EP4 axis

LECs are resistant to IR-related apoptosis and expand after IR exposure, where they serve as an essential source of pro-regenerative niche factors such as RSPO3, Wnt2a, and Ccl21a (Goto et al., 2022; Niec et al., 2022; Ogasawara et al., 2018; Palikuqi et al., 2022). Given this established role of lymphatics in intestinal regeneration, we sought to investigate if there could be crosstalk between LECs and Hdc^+^ IMCs after injury. Staining for LYVE1 in the intestines of Hdc^GFP^ mice 3 days after WB-IR revealed a close association between LECs and IMCs (**Figure 3A**).

**Figure 3:**
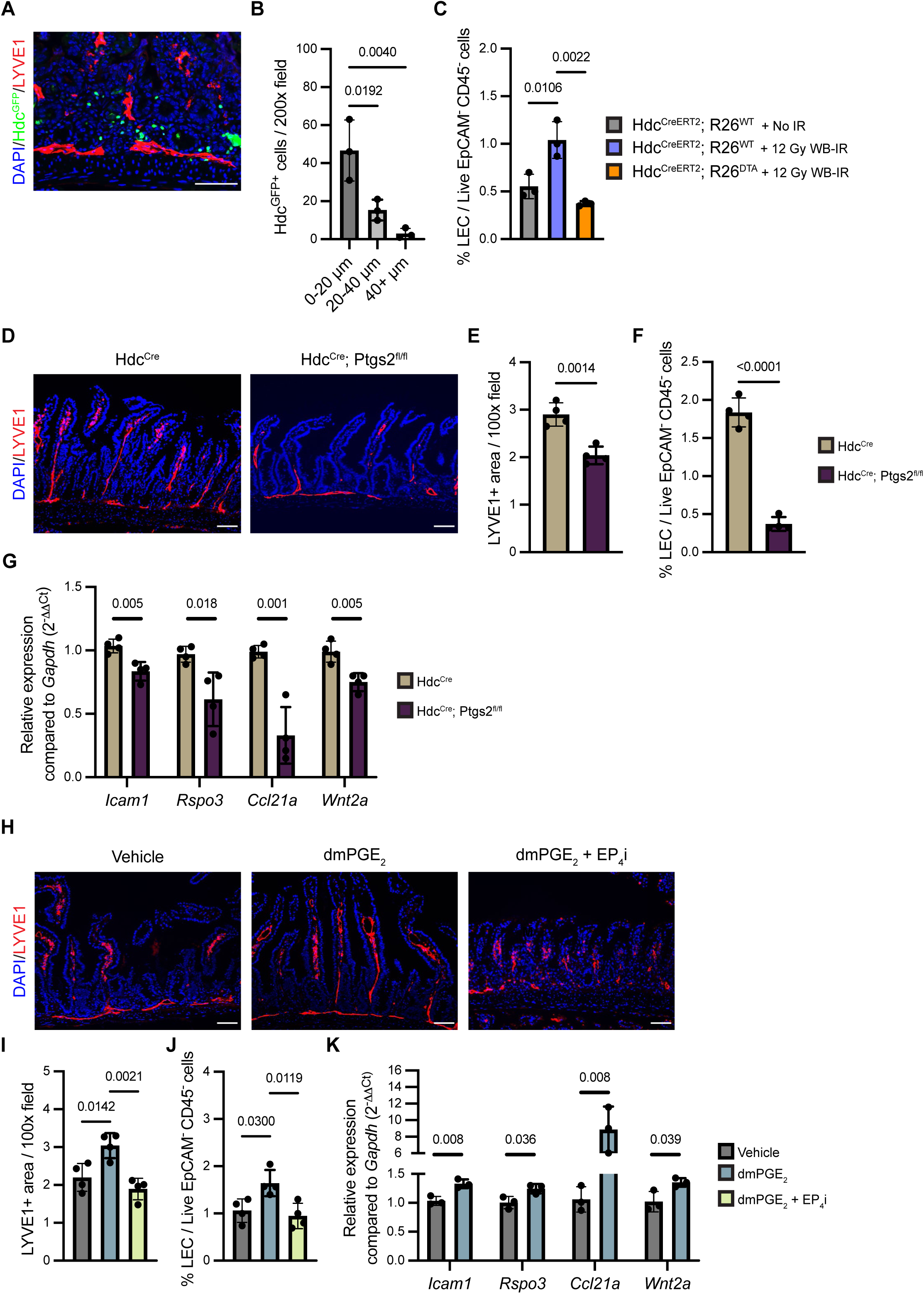
Hdc+ IMC-derived PGE2 activates intestinal lymphatics. **A.** Representative 200x image of LYVE1 staining with Hdc^GFP^ visualization from the proximal jejunum of Hdc^GFP^ mice after 12 Gy WB-IR. **B.** Bar plot showing quantification of the average number of GFP+ cells in within proximity ranges to LYVE1+ staining per 200x field. (n=3). **C.** Quantification of the percentage of CD90.2+ LECs (CD31^+^ CD90.2^+^) out of all DAPI^-^ CD45^-^ EpCAM^-^ cells from the small intestines of Hdc^CreERT2^; R26^DTA^ mice with no irradiation (No IR), Hdc^CreERT2^, R26^WT^ mice receiving 12 Gy WB-IR, and Hdc^CreERT2^; R26^DTA^ mice receiving 12 Gy WB-IR. See (Figure 1J) for experimental scheme. (n=3). **D.** Representative 100x images of LYVE1 staining from the proximal jejunum of Hdc^Cre^ or Hdc^Cre^; Ptgs2^fl/fl^ mice after 12 Gy WB-IR. See (Figure 2B) for experimental scheme. **E.** Bar plot showing quantification of the average LYVE1+ area per 100x field of images in (**J**). (n=4). **F.** Quantification of the percentage of CD90.2^+^ LECs (CD31^+^ CD90.2^+^) out of all DAPI^-^ CD45^-^ EpCAM^-^ cells from the small intestines of Hdc^Cre^ or Hdc^Cre^; Ptgs2^fl/fl^ after 12 Gy WB-IR. (n=4). **G.** qPCR analysis of *Icam1, Rspo3, Ccl21a,* and *Wnt2a* of sorted LECs from the intestines of Hdc^Cre^ or Hdc^Cre^; Ptgs2^fl/fl^ mice after 12 Gy WB-IR. (n=4). **H.** Representative 100x images of LYVE1 staining from the proximal jejunum of Hdc^GFP^ mice treated with vehicle, dmPGE2, or dmPGE2 with EP4i pretreatment after 12 Gy WB-IR. **I.** Bar plot showing quantification of the average LYVE1+ area per 100x field of images in (**H**). (n=4). **J.** Quantification of the percentage of CD90.2+ LECs (CD31^+^ CD90.2^+^) out of all DAPI^-^ CD45^-^ EpCAM^-^ cells from the small intestines of Hdc^GFP^ mice treated with vehicle, dmPGE2, or dmPGE2 + EP4i pretreatment after 12 Gy WB-IR. **K.** qPCR analysis of *Icam1*, *Rspo3, Ccl21a* and *Wnt2a* of sorted LECs from the intestines of mice treated with vehicle or dmPGE2 after 12 Gy WB-IR. Scale bars = 50 uM Bar graph data are mean ± SEM. Statistical analysis was performed using an Ordinary one-way ANOVA with multiple comparisons to each group (**B, C, I, J**) or a two-sided Student’s t test (**E, F, G, K**).

A population of RSPO3-secreting LECs in the intestine can be marked as CD31^+^ CD90.2^+^ (Ogasawara et al., 2018). We confirmed this population to be LECs by FACS sorting them then analyzing their expression of *Lyve1, Pdpn, Prox1,* and *Vegfr3* compared to sorted blood endothelial cells (CD31+ CD90.2-)(Murtomaki et al., 2013)(**Supplementary Figure S3A**). Ablation of Hdc+ cells using Hdc^CreERT2^; R26^DTA^ mice diminished this LEC population after IR injury compared to controls, while adoptive transfer of Hdc^GFP+^ IMCs (but not Hdc^GFP-^ IMCs) partially rescued this reduction (**Figure 3B, Supplementary Figure S3B**). Since our data indicate that Hdc^+^ IMC-derived PGE2 is important for regeneration, we suspected that the reduction of LECs in the absence of IMCs may be due to lack of IMC-derived PGE2. In support of this hypothesis, PGE2 has previously been described to induce lymphangiogenesis of LECs in vitro through the EP4 receptor (Nandi et al., 2017). We found that intestinal LECs isolated via FACS primarily express *Ptger4,* the gene encoding the EP4 receptor (**Supplementary Figure S3C**).

To determine if IMC-derived PGE2 is important for LEC activity during regeneration, we returned to the Hdc^Cre^; *Ptgs2*^fl/fl^ mice. After WB-IR, Hdc^Cre^; *Ptgs2*^fl/fl^ mice had diminished LYVE1+ area near the intestinal crypts compared to Hdc^Cre^ controls via immunostaining, and a smaller percentage of LECs via FACS analysis (**Figures 3C – E**). FACS-isolated LECs from Hdc^Cre^; *Ptgs2*^fl/fl^ mice also had reduced expression of the activation marker *Icam1*, and the pro-regenerative genes *Rspo3, Ccl21a,* and *Wnt2a* compared to controls (Niec et al., 2022)(**Figure 3F**). Together, these data indicate that Hdc^+^ IMC-derived PGE2 is important for activating LECs to expand and upregulate regenerative factors after IR injury.

To further investigate the effects of PGE2 on LECs, we performed bulk RNA sequencing on human dermal lymphatic endothelial cells (HDLEC) treated with dmPGE2. Analysis of the pathways that are upregulated by dmPGE2 treatment revealed genes involved in endothelial cell adhesion, migration, wound healing, inflammatory response, and angiogenesis (**Supplementary Figure S3D**). To confirm if dmPGE2 can induce lymphangiogenesis of HDLECs, we performed an *in vitro* endothelial cell bead sprouting assay (Azam et al., 2018). HDLECs sprouted when treated with dmPGE2, but this was inhibited by EP4i treatment (**Supplementary Figures S3E – S3F**). Additionally, dmPGE2-treated HDLECs had higher expression of activation and regenerative factors, and RSPO3 secretion, compared to cells treated with a vehicle, but this effect could be diminished by co-treating the cells with an EP4 antagonist (EP4i) (**Supplementary Figures S3G – H**). These results indicate that PGE2 promotes LEC activation, sprouting, and expression of pro-regenerative factors through the EP4 receptor.

Given the importance of IMC-derived PGE2 during intestinal regeneration, we then sought to determine if dmPGE2 treatment could induce additional lymphangiogenesis in the intestines of irradiated mice and thus improve regeneration. Mice that received dmPGE2 after 12 Gy WB-IR had increased LEC percentage as well as lymphatic expansion near the SI crypt bases compared to controls, as shown by LYVE1 immunostaining and FACS analysis, and this effect was blocked by EP4i pretreatment (**Figures 3G – 3I**). FACS-isolated intestinal LECs from mice treated with dmPGE2 showed higher gene expression of activation and pro-regenerative factors (**Figure 3J**). Additionally, dmPGE2 injected into unirradiated mice also increased LEC expression of activation and regenerative factors independent of barrier damage (**Supplementary Figure S3I**).

Taken together, these data indicate that PGE2 from infiltrating Hdc^+^ IMCs, besides promoting epithelial proliferation, also act upon LECs via the EP4 receptor to promote activation, sprouting, and expression of pro-regenerative factors.

### Hdc+ IMCs are recruited to the damaged intestine via CXCR2 and CXCR4 signaling

To better understand the connection between intestinal damage, Hdc^+^ IMCs, and LECs, we then investigated how Hdc^+^ IMCs are initially recruited to the damaged tissue. To this aim, we performed the RT^2^ Profiler PCR array for chemokine receptors on Hdc^GFP+^ cells isolated from intestines of mice 3 days after WB-IR. Results showed that these cells express high levels of the chemokine receptor genes *Cxcr2* and *Cxcr4*, consistent with previous literature on granulocytic IMCs (Katoh et al., 2013) (**Figure 4A**). To determine if CXCR2 and CXCR4 signaling are important for Hdc+ IMC recruitment to the damaged intestine, we treated Hdc^GFP^ mice with a CXCR2 or CXCR4 inhibitor before and after exposure to IR (**Figure 4B**). Inhibition of CXCR2 or CXCR4 was sufficient to significantly reduce the number of Hdc^+^ cells infiltrating the intestine via histological and FACs analysis of GFP^+^ cells, suggesting that these receptors are important for IMC trafficking to the IR-injured intestine (**Figures 4C – E**).

**Figure 4:**
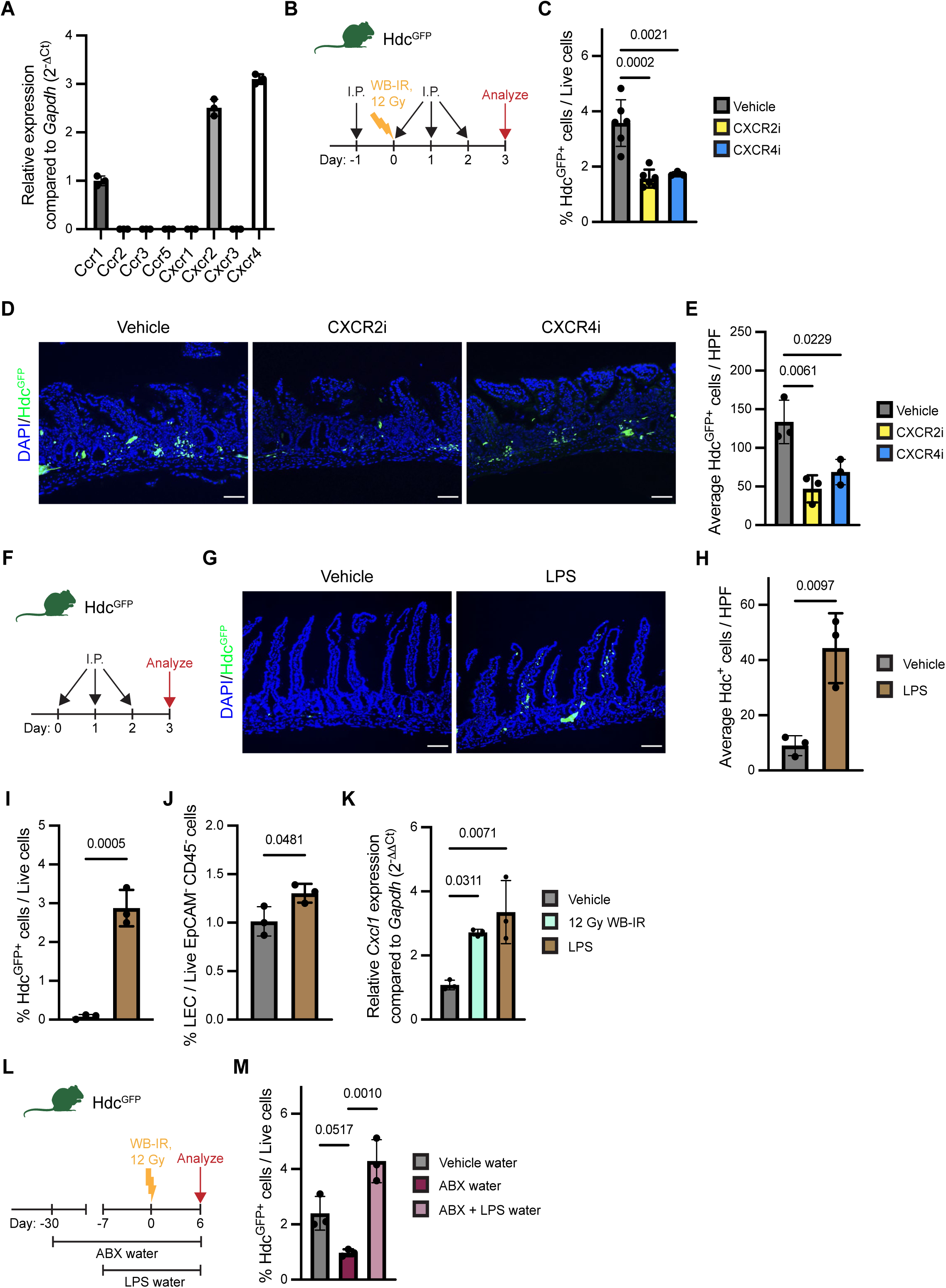
Microbial signals modulate the crosstalk between immature myeloid cells and lymphatics. **A.** RT^2^ gene expression analysis of chemokine receptor genes of sorted Hdc^GFP+^ IMCs from the intestine of mice after 12 Gy WB-IR. (n=3). **B.** Experimental scheme of CXCR2/CXCR4 inhibition experiment. I.P.= intraperitoneal injection of vehicle, SB225002 (CXCR2i) or MSX-122 (CXCR4i). **C.** Bar plot showing quantification of FACS analysis of the percentage of Hdc^GFP+^ cells out of live (DAPI-) cells from the small intestines of Hdc^GFP^ mice treated with vehicle, CXCR2i, or CXCR4i before and after 12 Gy WB-IR. **D.** Representative 100x images of GFP visualization from the proximal jejunum of Hdc^GFP^ mice that received vehicle, CXCR2i, or CXCR4i before and after 12 Gy WB-IR. **E.** Bar plot showing quantification of GFP+ cells of images in (**D**). (n=3), HPF: high powered field. **F.** Experimental scheme of LPS injection experiment. I.P. = intraperitoneal injection of vehicle or LPS. **G.** Representative 100x images of GFP visualization from the proximal jejunum of Hdc^GFP^ mice that received vehicle or LPS treatment. **H.** Bar plot showing quantification of Hdc^GFP+^ cells of images in (**G**). (n=3), HPF: high powered field. **I.** Bar plot showing quantification of FACS analysis of the percentage of Hdc^GFP+^ cells out of live (DAPI^-^) cells from the small intestine of Hdc^GFP^ mice after vehicle or LPS injections. **J.** Quantification of the percentage of CD90^+^ LECs (CD31^+^ CD90.2^+^) out of all DAPI^-^ CD45^-^ EpCAM^-^ cells from the small intestines of Hdc^GFP^ mice after vehicle or LPS injections. (n=3). **K.** qPCR analysis of *Cxcl1* of sorted LECs from the intestines of mice treated with vehicle, 12 Gy WB-IR, or LPS injections. (n=3). **L.** Experimental scheme of microbiome knockdown experiment. ABX = antibiotic cocktail. **M.** Bar plot showing quantification of FACS analysis of the percentage of Hdc^GFP+^ cells out of live (DAPI^-^) cells from the small intestine of mice after irradiation given different drinking waters. (n=3). Scale bars = 50 uM Bar graph data are mean ± SEM. Statistical analysis was performed using an Ordinary one-way ANOVA with multiple comparisons to each group (**C, E, K, M**) or a two-sided Student’s t test (**H, I, J**).

### The microbiome modulates Hdc+ IMC recruitment and LEC expansion

Next, we asked how CXCR2 and CXCR4 signaling could be initiated during intestinal injury. During intestinal barrier disruption, commensal microbes that reside in the lumen breach the barrier and contact the epithelium and underlying mesenchyme (Nighot et al., 2017). Lipopolysaccharides (LPS) are a major component of the cell walls of gram-negative bacteria, and mediate intestinal inflammatory responses (Guo et al., 2015). Previous studies have reported that exposing lymph node LECs to LPS causes them to secrete CXCL1, a chemokine that can bind to CXCR2 (Xu et al., 2012). We therefore asked if LPS exposure could serve as a trigger for IMC recruitment via LECs into the intestine. To test this, we injected healthy Hdc^GFP^ mice with LPS then analyzed the abundance of Hdc^GFP+^ cells in the intestine (**Figure 4F**). Within 3 days, significantly more Hdc^+^ IMCs infiltrated the intestinal mucosa of LPS-treated mice compared to controls (**Figures 4G – I**). This was associated with an increased LEC percentage (**Figure 4J**). Notably, intestinal FACS-isolated LECs upregulated *Cxcl1* when exposed to IR or LPS *in vivo* (**Figure 4K**). We validated these findings using an *in vitro* LEC model in which HDLECs were stimulated with LPS, and then showed significant upregulation of *Cxcl1* and *Cxcl8* (encoding IL-8, the human-specific CXCR2 ligand) upon LPS exposure (**Supplementary Figures S4A**). Additionally, LPS-stimulated HDLECs actively promoted migration of HDC+ IMCs, but this migration could be abrogated using a CXCR2 inhibitor or the MEK1/2 antagonist U0126, which has been shown to inhibit CXCR2 signaling (Yang et al., 2019) (**Supplementary Figures S4B – C**).

Next, to test if LPS signals coming from gut microbes are important for recruiting IMCs, we treated Hdc^GFP^ mice with an antibiotic cocktail in their drinking water for 30 days to deplete their microbiome, then challenged them with WB-IR (**Figure 4L**). After IR injury, antibiotic-treated mice had significantly fewer Hdc^GFP+^ cells infiltrating the intestine, highlighting the importance of microbial invasion in this pathway (**Figure 4M**). Addition of LPS to the water of the antibiotic-treated mice with LPS 7 days before irradiation rescued the recruitment of Hdc^GFP+^ cells into the intestine. These results suggest that signals coming from gut microbes activate LECs to promote IMC infiltration and initiate intestinal repair after epithelial barrier disruption.

## DISCUSSION

The mammalian intestine has evolved symbiotically with the gut microbiome and has developed mechanisms of interactions with these microbes to modulate host metabolism, immunity, and physiology. Breaches of the intestinal barrier cause disruptions to the balance of normal microbial interactions, requiring a response from the host to restore homeostasis. LECs have been established as key niche cells for the ISC, as they expand during injury and are a major source of RSPO3 (Goto et al., 2022; Niec et al., 2022; Ogasawara et al., 2018; Palikuqi et al., 2022). While LECs have been shown to be central to the intestinal regeneration program, the mechanisms leading to their activation following injury were not known. Here, we have shown a crucial role for enteric microbes and immature myeloid cells in regulating intestinal regeneration, in part through modulating LEC activity.

Understanding the mechanism of mucosal repair is crucial for improving therapy for tissue injury. Until now, the involvement of IMCs as a regenerative cell type has been understudied. We report here that Hdc^+^ IMCs are rapidly recruited to the IR-injured intestine, where they are necessary for proper regeneration. Moreover, we found that the regenerative effects of Hdc+ IMCs can be attributed, at least in part, to their secretion of PGE2. In line with previous reports, we found that IMC-derived PGE2 exerts direct mitogenic effects on intestinal epithelial cells (Miyoshi et al., 2017; Roulis et al., 2020). Furthermore, we found that PGE2 can act on LECs via EP4 to stimulate lymphangiogenesis and upregulate the expression of pro-regenerative genes, such as *Rspo3*, *Ccl21a*, and *Wnt2a*.

IMCs are typically mobilized in response to infection, stress, and tumorigenesis, but have been shown by us to function as key bone marrow niche cells, promoting the quiescence and self-renewal of long-term hematopoietic stem cells (HSCs) (Chen et al., 2017). As precursors for neutrophils and monocytes, IMCs are present at low levels outside of the bone marrow during homeostasis but become more abundant during disease states. These cells are rapidly mobilized in response to inflammatory conditions to fight infection and promote tissue repair, but such mobilization promotes activation of HSCs, which can eventually lead to their exhaustion if such activation is sustained (Fu et al., 2021). Although it was known these cells infiltrate damaged tissues to promote regeneration, their functions therein have been understudied. Here, we show that Hdc+ IMCs can be recruited by activated LECs, resulting in signals for LEC expansion and secretion of regenerative factors that promote regeneration of the epithelium.

Besides RSPO3, LECs also secrete other angiocrine factors that support stem cells and regenerative processes throughout the body (Augustin and Koh, 2017; Palikuqi et al., 2020; Rafii et al., 2016). Our findings show one pathway in which LECs are activated to promote tissue regeneration, as mediated by microbial signals and IMCs. The conclusion that IMCs can promote lymphangiogenesis have wider implications than in just regeneration. Lymphangiogenesis has been shown to be involved the development of breast cancer, colon cancer metastasis, and inflammatory bowel disease (Nandi et al., 2017; Ocansey et al., 2021; Tacconi et al., 2015). Thus, given their role in promoting LEC activation and sprouting by producing PGE2, it may be worth investigating the role of IMCs in these other conditions.

Intriguingly, we found that the recruitment of Hdc^+^ IMCs to the intestine is initiated by exposure of intestinal lymphatics to invading microbial signals. Specifically, LPS triggers LECs to upregulate *Cxcl1*, the ligand for CXCR2, which is highly expressed on Hdc^+^ IMCs. Treatment of healthy mice with LPS reproduced this recruitment pathway, and depletion of the intestinal microbiome abrogated recruitment following IR injury, confirming the key role of microbes in triggering Hdc^+^ IMC migration. The finding that enhanced penetration of gut microbes after mucosal injury is the initiating signal for IMC recruitment, LEC activation and intestinal regeneration suggests that loss of barrier integrity is not only pathogenic but also regenerative. Indeed, while severe injury and bacterial infiltration may lead to sepsis, moderate levels of microbial exposure appear to promote regenerative signals needed to restore intestinal health and homeostasis. Thus, at least in the intestine, microbial signals serve as physiologic signals that can initiate mucosal repair.

### Limitations of the study

In our work, we showed that Hdc+ IMCs are indispensable for proper SI repair after WB-IR. It has been previously reported that after exposure to radiation, the bone marrow is suppressed and may fail at higher doses (Green and Rubin, 2014). Given that IMCs originate from the bone marrow, it is possible that some of the effects we observed may be due to alterations to the bone marrow after IR. Thus, the WB-IR model may not fully reflect the circumstances of other types of intestinal injury, and more work should be done to observe the role of IMCs in other models, such as abdominal irradiation (AIR) with bone marrow shielding.

## ACKNOWLEDGEMENTS

This research was funded in part through the NIH/NCI Cancer Center Support Grant P30CA013696 and used the resources of the Herbert Irving Comprehensive Cancer Center Flow Cytometry Shard Resources, Molecular Pathology/MPSR, and Genomics and High Throughput Screening. This work was supported by grants from the NIH/NCI including R01DK48077 and R35CA210088 to T.C.W.; as well as an NCI Outstanding Investigator Award (R35 CA197745). Q.W. was supported in part by the Columbia University Institute of Human Nutrition’s training grant (T32 DK007647-32). This research was funded in part through the NIH/NCI Cancer Center Support Grant P30CA013696 and used the Genomics and High Throughput Screening Shared Resource. We thank Harvey Herschman for the generous gift of the Ptgs2^fl/fl^ mice. We thank Leah Zamechek, Jonathan LaBella, and Harry Nagendra for their support of our animal colony. We are grateful for the support we received by Columbia University shared resources, and we would like to thank Sun Dajiang “Kevin” (Molecular Pathology/MPSR), Michael Kissner (Columbia Stem Cell Initiative Flow Cytometry Core Facility), and Erin Bush (Genomics and High Throughput Screening) for their expertise and help during this project.

## AUTHOR CONTRIBUTIONS

Conceptualization, Z.J., Q.T.W, and T.C.W.; Methodology, Z.J., Q.T.W., W.K., E.M., C.G., C.S., and K.Y.; Investigation, Z.J., Q.T.W, N.F., and W.K.; Writing – Original Draft, Z.J. and Q.T.W.; Writing – Review & Editing, Z.J., Q.T.W., E.M., C.G., C.S., K.Y., and T.C.W.; Funding Acquisition, T.C.W.; Supervision, C.G., C.S., K.Y., and T.C.W.

## STAR METHODS

### KEY RESOURCES TABLE

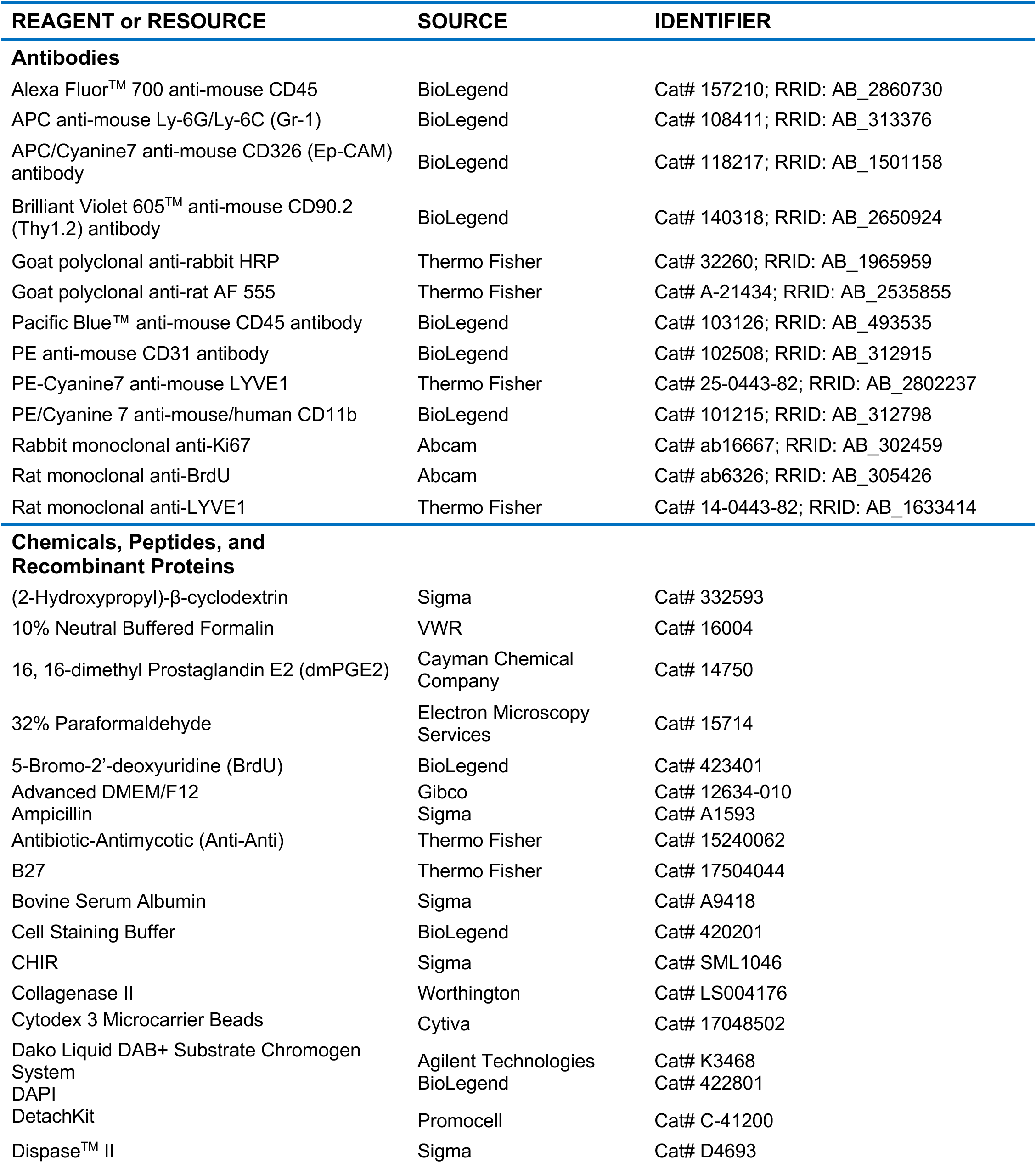
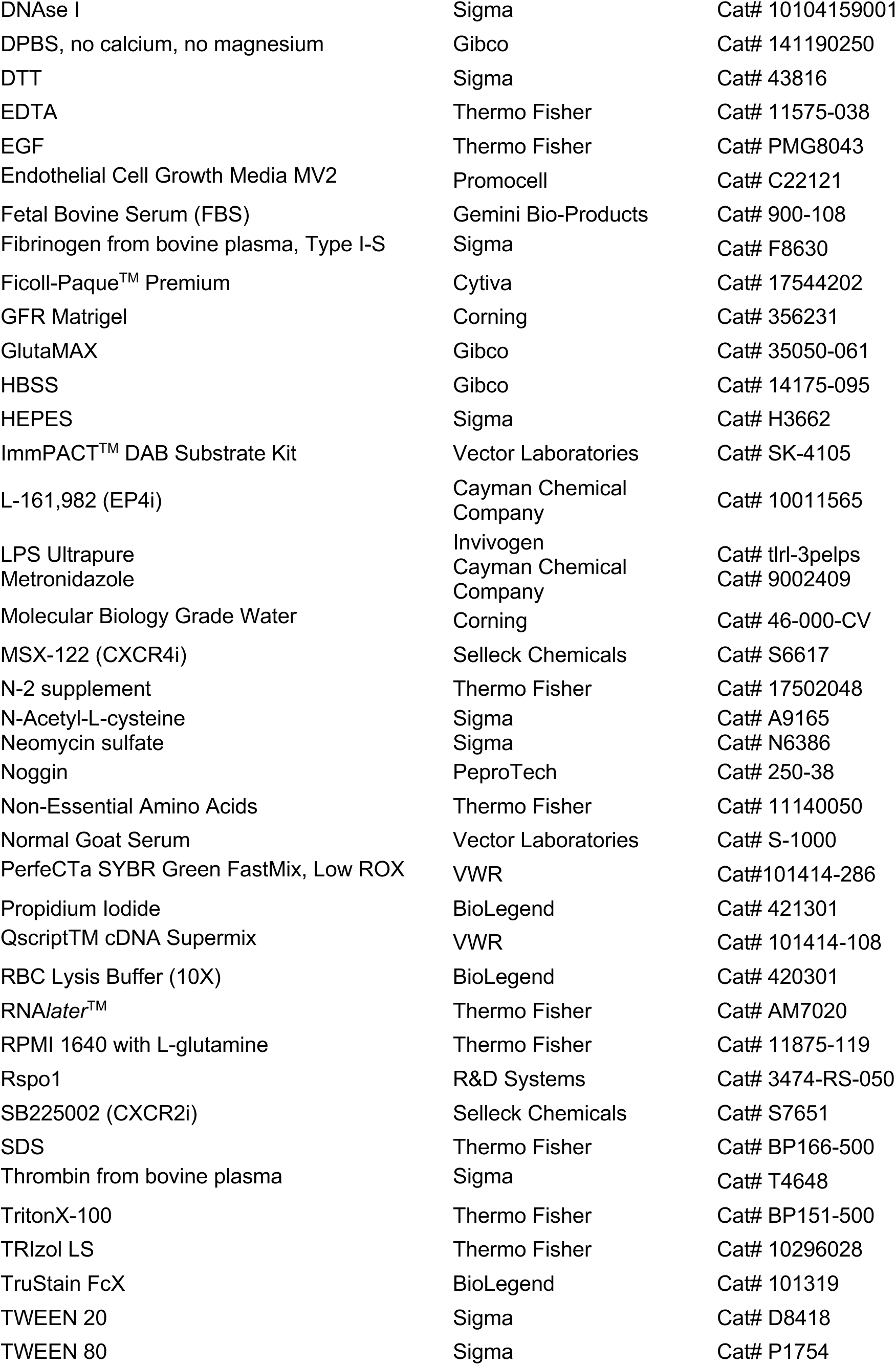
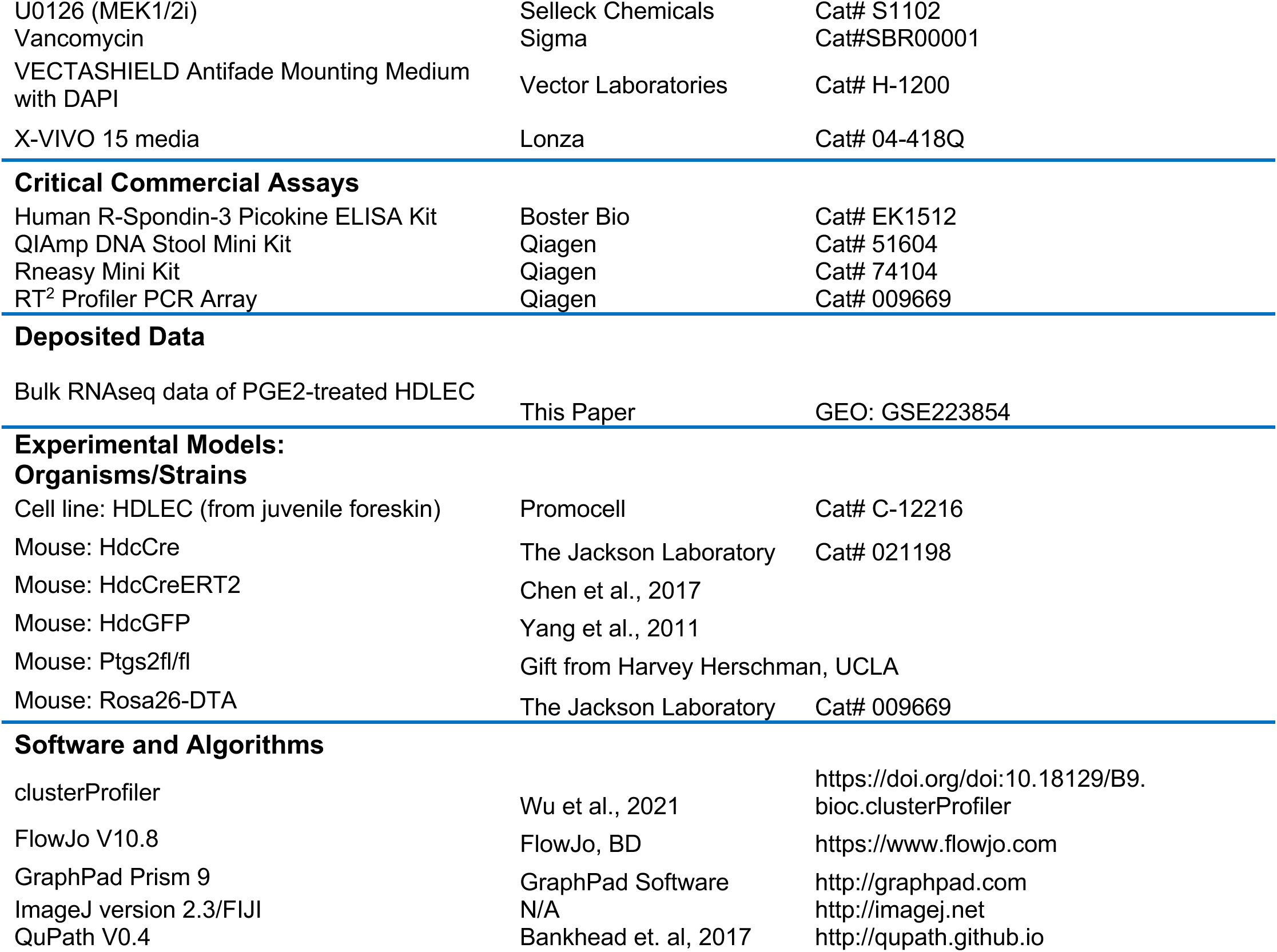

### CONTACT FOR REAGENT AND RESOURCE SHARING

Requests for resources and reagents should be directed to and will be fulfilled by the Lead Contact, Timothy C. Wang.

### EXPERIMENTAL MODEL AND SUBJECT DETAILS

#### Mouse crosses and care

All mouse experiments were conducted under the IACUC protocol AABE1556 at the Columbia University Irving Medical Center, and all mouse studies were approved by the Columbia University Institutional Animal Care and Use Committee. All mice (male and female) were used at 8-12 weeks of age at the beginning of each experiment. *Hdc^GFP^* and *Hdc^CreERT2^*mice have been described previously (Chen et al., 2017; Yang et al., 2011). *Hdc^CreERT2^* was mated to *Rosa26-DTA* (Jackson labs 007909) (R26^DTA^) mice for generating *Hdc^CreERT2^; R26^DTA^* mice. *Hdc^Cre^*mice were purchased from Jackson labs (021198 (Zecharia et al., 2012). *Ptgs2^fl/fl^* mice were gifted by Harvey Herschman (Ishikawa and Herschman, 2006). *Hdc^Cre^* was mated to *Ptgs2^fl/fl^* to generate *Hdc^Cre^; Ptgs2^fl/fl^*mice. These mice were maintained in a C57BL/6 background.

#### Mouse drug treatments

Mice were given the CXCR2 inhibitor SB225002 (Selleck Chemicals, S7651) at a dose of 1 mg/kg in PBS + 0.33% Tween 80, or vehicle, intraperitoneally daily for 4 consecutive days. The CXCR4 inhibitor MSX-122 (Selleck Chemicals, S6617) was administered at a dose of 10 mg/kg in PBS +10% DMSO +45% (2-Hydroxypropyl)-β-cyclodextrin (Sigma, 332593) vehicle, or vehicle, intraperitoneally daily for 4 consecutive days. 16,16-dimethyl Prostaglandin E2 (dmPGE_2_, Cayman Chemical Company, 14750) was administered intraperitoneally at a dose of 1 mg/kg in PBS +10% ethanol, or just this vehicle, daily for 3 consecutive days. The EP_4_ receptor inhibitor L-161,982 (Cayman Chemical Company, 10011565) was administered intraperitoneally at a dose of 10 mg/kg in PBS +10% ethanol one hour before dmPGE_2_ administration. For LPS injections, LPS (Invivogen, tlrl-3pelps) was dissolved in PBS, and this or just PBS was injected intraperitoneally at a dose of 2mg/kg. For LPS treatment in drinking water, LPS was dissolved in sterile water (10 ug/mL) and given to mice for 13 days. For Hdc+ cell ablation and Cre activation, mice were fed a tamoxifen diet (delivering 80 mg/kg body weight daily, Envigo TD.130858) for 2 days before irradiation, and continued the diet until the experiment’s end. For BrdU proliferation experiments, 100 μL of BrdU (10 mg/mL, BioLegend 423401) was injected intraperitoneally in the mice 2 hours before euthanasia.

#### Irradiation experiments

Mice were challenged with 12 Gy WB-IR delivered by either a cesium 137 irradiator or a Multirad 350 X-Ray irradiator (Precision X-Ray). Tissue was collected at various days post-irradiation.

#### Antibiotic treatments

Mice were given an antibiotic cocktail diluted in their drinking water for 30 days. The cocktail consisted of ampicillin (1 g/L) (Sigma, A1593), neomycin sulfate (1 g/L) (Sigma, N6386), metronidazole (1 g/L) (Cayman Chemical Company, 9002409), and vancomycin hydrochloride (0.5 g/L) (Sigma, SBR00001). Bacterial depletion was measured using the QIAmp DNA stool mini kit and following manufacturer instructions.

### METHOD DETAILS

#### Tissue collection and preparation for microscopy

For all microscopy experiments, the small intestine was collected, cleaned with PBS, then fixed in 4% PFA or 10% neutral-buffered formalin (VWR 16004) at 4°C for 24 hours. For sectioning, fixed tissues were embedded in paraffin or OCT before being sectioned to a thickness of 5 μm.

#### Immunohistochemistry, immunofluorescence and BrdU labeling

For immunohistochemical staining, formalin-fixed, paraffin-embedded (FFPE) sections were deparaffinized in xylene and rehydrated in gradient ethanol. Antigen retrieval was performed by heating the slides in citrate buffer in a steamer for 40 min. Endogenous peroxidase was blocked by incubation with 3% hydrogen peroxide (Sigma) in PBS. Slides were rinsed, blocked with 10% serum, and incubated with primary antibodies (anti-Ki67, Abcam, ab16667) overnight at 4 °C. Subsequently, slides were incubated with secondary antibodies (diluted in 2% BSA) at RT for 45 min followed by peroxidase-conjugated avidin (Vector Laboratories) and 3,3’-diaminobenzidine (Dako) as a chromogen. Slides were counterstained with hematoxylin.

Frozen slides were permeabilized with 1% SDS in PBS at room temperature for 5 minutes, blocked with 10% normal goat serum in 0.1% Tween 20 at room temperature for 1 hour, and stained with primary antibody (rat monoclonal anti-LYVE1, 1:1000, Thermo Fisher Scientific, 14-0443-82) at 4°C overnight. Then, sections were incubated with secondary antibodies at room temperature for 2 hours (goat polyclonal anti-Rat IgG conjugated to AF555, 1:400, Thermo Fisher Scientific, A-21434). All slides were counterstained and mounted with VECTASHIELD Antiface Mounting Medium with DAPI (Vector Laboratories). Fluorescent images were acquired with an X-Cite Series 120 illuminator (Excelitas) on a Nikon Eclipse TE2000-IJ microscope stand (Nikon Instruments). H&E images were acquired with an AT2 microscope slide scanner (Leica) then processed using QuPath v0.4 (Bankhead et al., 2017).

For *BrdU* fluorescent staining, slides were incubated with 10 mg/mL DNAse I in PBS at 37°C for 20 minutes. Then, BrdU staining was performed as above using an anti-BrdU antibody (rat monoclonal anti-BrdU, 1:100, Abcam ab6326).

#### Quantification of staining

All image analysis was done using Fiji software (Schindelin et al., 2012). The average number of Hdc^GFP+^ cells per high-powered field (HPF) was quantified by counting the number of Hdc^GFP+^ cells for 5 randomly selected image fields per mouse, then averaged. The average number of BrdU+ cells per crypt was quantified by counting the number of BrdU+ cells per crypt for every crypt in 5 separate 100x image fields, then calculating the mean for each mouse. For the mean villus length, the length of every villus was measured in 5 separate 100x image fields per mouse then averaged. Lymphatic vessel density was quantified by measuring the average area of LYVE-1 positive area on total area for 5 separate 100x image fields per mouse. Images are all composites of different channels unless otherwise stated.

#### Small intestine tissue digestion and flow cytometry analysis

Lymphatic cells were isolated from the small intestine as previously described (Couter and Surana, 2016; Palikuqi et al., 2022). Briefly, the small intestine tissues from adult male and female mice were collected, flushed with cold PBS and dissected to remove mesentery and fat tissue. Then, the tissue was cut into 3 cm pieces, canaliculated, and the mucus was removed with a paper towel. The pieces were put into 30 mL of dissociation solution (RPMI 1640 with L-glutamine (Thermo Fisher 11875-119) + 1% non-essential amino acids (Thermo Fisher 11140050), 2.5% HEPES (Sigma H3662), 1% Antibiotic-Antimycotic (Thermo Fisher 15240062), 5 mM EDTA (Thermo Fisher 11575-038), 3% FBS (Gemini Bio-Products 900-108) and 1 μM DTT (Sigma 43816), and incubated on a rotator at 37°C for 10 minutes. Then, the tissues were strained through a 100 μM cell strainer, washed twice with HBSS, and minced finely using scissors. The minced tissue was then placed in 4 mL of digestion media (HBSS + 1% Glutamax, 1% non-essential amino acids, and 2.5% HEPES with 2.5 mg/mL Collagenase II (Worthington LS004176), 1 U/mL Dispase II (Sigma D4693), and 30 μg/mL DNAse I (Sigma 10104159001)) and incubated at 37°C for 15 minutes on a rotator. The solution was passed through a 14-gauge needle 5 times using a syringe, then placed back on the rotator to digest for 5 more minutes at 37°C. The digested cells were strained through a 100 μM strainer, pelleted, then placed in 5 mL 1x RBC lysis buffer (BioLegend 420301) for 5 minutes on ice. The reaction was quenched with PBS, then spun down, and the remaining single cell suspension was incubated with TruStain FcX (BioLegend, 101319) for 5 minutes on ice. Finally, the suspension was stained with the following antibody cocktail: CD45-Pacific Blue (1:200, BioLegend, 103125), TER119-Pacific Blue (1:200, BioLegend, 116231), Ep-CAM-APC-Cy7 (1:200, BioLegend, 118217), CD31-PE (1:500, BioLegend, 160203), Lyve1-PE-CY7 (1:300, eBioscience, 25-0443-82), CD90.2-BV605 (1:300, BioLegend, 105343). Propidium iodide (BioLegend 421301) or DAPI (BioLegend, 422801) was used as a viability dye to exclude dead cells from the analysis. Cells were analyzed on a NovoCyte Quanteon or sorted using a Sony MA900 with a 100-micron nozzle. GFP+ cells were defined as PI^-^, GFP^+^. Lymphatic endothelial cells were defined as PI^-^, Ep-CAM^-^, CD45^-^, TER119^-^, CD31^+^, CD90.2^+^, Lyve1^+^. For RNA isolation, cells were sorted directly into TRIzol LS (Thermo Fisher 10296028). The data was analyzed using FlowJo software (version 10, TreeStar).

#### Bone marrow cell isolation

HDC^+^ bone marrow cells were obtained from Hdc^GFP^ mice by crushing leg, arm, and pelvic bones in HBSS containing 2% heat-inactivated fetal bovine serum (Gemini, 900-108). Single-cell suspensions were purified on a Ficoll gradient (Cytiva, 17544202), then red blood cells were lysed using 1X RBC lysis buffer. The cells were stained with the following antibody cocktail: CD45-AF700 (1:800, BioLegend, 103127), CD11b-PE-Cy7 (1:800, BioLegend, 101215), Gr-1-APC (1:400, BioLegend, 108411). Propidium iodide was used as a viability dye to exclude dead cells. Cells were sorted using a Sony MA900 with a 100-micron nozzle. CD45^+^, CD11b^+^, Gr-1^+^, GFP^+^ cells were sorted directly into X-VIVO 15 media (Lonza, 04-418Q) for further culture experiments, or directly into TRIzol LS for RNA isolation.

#### Adoptive transfer

HdcCre^ERT2^; R26^DTA^ mice were placed on a tamoxifen diet for 2 days, then were challenged with 12 Gy WB-IR. CD11b^+^ Hdc^GFP+^ or CD11b^+^ Hdc^GFP-^ myeloid cells were isolated from the bone marrow of several Hdc^GFP^ donor mice and pooled. 6 hours after WB-IR, HdcCre^ERT2^; R26^DTA^ recipient mice were injected with 3 million CD11b^+^ Hdc^GFP+^ or CD11b^+^ Hdc^GFP-^ cells via the tail vein, and this was repeated 72 hours later. Samples were collected 6 days after WB-IR.

#### *In vitro* organoids

The small intestine from adult male and female mice were collected, flushed with cold PBS, cleaned of the mesentery, and opened lengthwise. EDTA (Invitrogen, 15575020) based dissociation was performed as previously described (Sato et al., 2009). Following EDTA dissociation crypts were resuspended in Advanced DMEM/F12 (Gibco, 12634-010) supplemented with GlutMAX (Thermo Fisher Scientific, 35050061), HEPES (Thermo Fisher Scientific, 15-630-080), Antibiotic-Antimycotic (Anti/Anti - Fisher Scientific, 15240062), 10% FBS (Gemini Bio-Products, 900-108), B27 (Thermo Fisher Scientific, 17504044), N-2 supplement (Fisher Scientific, 17502048), N-Acetyl-L-cysteine (Sigma-Aldrich, A9165), EGF (Thermo Fisher Scientific, PMG8043), Noggin (PeproTech, 250-38), Rspo1 (R&D Systems, 3474-RS-050), and CHIR (Sigma-Aldrich, SML1046). Crypts were counted under a microscope, then 200 crypts were seeded into 20 uL GFR Matrigel (Corning 356231) and plated in a pre-warmed 24-well plate as domes. dmPGE2 and/or EP4i treatment began 24 hours after treatment, and media was replaced every two days. Organoid size and number data was collected on day 6 after plating.

#### HDLEC cell culture

Human dermal lymphatic endothelial cells (HDLEC) from juvenile foreskin were purchased from Promocell (C-12216) and cultured in Endothelial Cell Growth Media MV2 (Promocell C22121). Cells were maintained in a humidified incubator set to 37°C and 5% CO_2_. For PGE_2_ stimulation experiments, 3x10^5^ cells per well were plated in a 12-well plate and allowed to seed overnight. dmPGE_2_ was added to a concentration of 500 nM or 1 μM, and L-161,982 was added to a concentration of 2 μM. For gene expression analysis, the cells were treated for 6 hours then lysed with Buffer RLT Plus, then RNA was collected using the RNEasy Plus Mini Kit (Qiagen 74134). For quantification of supernatant RSPO3, the cells were treated for 18 hours then the supernatant was harvested for ELISA analysis. For LPS stimulation, 3x10^5^ cells per well were plated in a 12-well plate and allowed to seed overnight. Ultrapure LPS (Invivogen, tlrl-3pelps) was added at a concentration of 1, 10, or 100 ng/mL and the cells were incubated for 6 hours before being harvested for RNA isolation or migration analysis.

#### HDLEC sprouting assay

An endothelial cell assay was done as described previously (Azam et al., 2018). Briefly, Cytodex 3 microcarrier beads (Cytiva 17048502) were coated by adding 10^6^ detached HDLEC to 1200 beads suspended in PBS. The bead-cell suspension was incubated at 37°C for 4 hours, agitating every 20 minutes. Then, the coated beads were plated on a 6 cm plate in Endothelial Cell Growth Media MV2 overnight. The next morning, the beads were inspected under a microscope to ensure consistent coating of the beads. The beads were washed then resuspended in a fibrinogen solution (Sigma, F8630), then plated into thrombin (Sigma, T4648) to form a fibrin gel in a 24-well glass-bottomed plate. Once the gel was fully solidified, the cells were treated with control media or media containing dmPGE_2_ at a concentration of 1 μM with or without L-161,982 at a concentration of 2 μM. The media was changed after 24 hours, then the cells were inspected for sprouting under a microscope 48 hours after implantation into the gel. Average sprout length per group was calculated by measuring the length of every sprout coming off 5 different beads per well, then calculating the mean.

#### HDC^+^ cell and HDLEC co-culture and migration assays

For migration analyses, HDLEC that had been stimulated with 0, 1, 10, or 100 ng/mL LPS were detached using a DetachKit (PromoCell, C-41200), pelleted, then washed twice with PBS. 5x10^4^ cells were plated in the bottom well of a 96-well Transwell plate with 5.0 μm pores (Corning, 3387), and allowed to seed overnight. The CXCR2 inhibitor SB225002 was added to some of the HDLEC-containing wells at a concentration of 200 nM, or the MEK1/2 inhibitor U0126 (Selleck Chemicals, S1102) was added to some wells at a concentration of 5 μM. 10^4^ sorted HDC^+^ immature myeloid bone marrow cells from Hdc^GFP^ mice were placed in the upper insert of the Transwell, and the plate was incubated for 12 hours. The top insert was discarded, and the average number of migratory cells per 200x field was calculated by counting the number of GFP^+^ cells for 5 independent 200x fields per bottom well then calculating the average.

#### Enzyme-linked immunosorbent assay (ELISA)

The concentration of RSPO3 was measured using the human R-Spondin-3 Picokine ELISA Kit (Boster Bio, EK1512) according to manufacturer’s instructions. HDLEC-conditioned media (18hr post treatment with vehicle or dmPGE2 with or without L-161,982) was used at 1, 1:2, and 1:4 serial dilutions. Protein concentration was measured at 450 nm on a SpectraMax iD3 Microplate Reader (Molecular Devices).

#### RNA isolation and cDNA synthesis

RNA was isolated from fresh whole tissue, tissue that had been preserved in RNAlater (Invitrogen, AM7020), or cell pellets. 5-10 mg of preserved distal small intestinal tissue from control or irradiated Hdc^GFP^ mice treated with dmPGE_2_ with or without L-161,982 was homogenized using metal beads in a Bullet Blender Tissue Homogenizer (Next Advance). The resulting homogenate was dissolved in Buffer RLT Plus (Qiagen) and RNA was isolated using the RNEasy Plus Micro kit (Qiagen, 74030) according to manufacturer’s instructions, including gDNA elimination. The final elution volume was 14 or 30 μL and RNA quantity and quality were measured on a NanoDrop 8000 spectrophotometer (Thermo Fisher). For RNA isolation from sorted bone marrow or small intestinal lymphatic endothelial cells, 1-5x10^5^ cells were sorted directly into TRIzol LS. RNA was then isolated via chloroform phase separation and isopropanol precipitation. RNA from cultured HDLEC was collected by lysing 80-90% confluent cells seeded in a 12-well plate in 500 mL Buffer RLT Plus, then homogenized using a QIAshredder column. The same amount of RNA (up to 1 ug) was loaded per sample for subsequent cDNA synthesis. cDNA was synthesized by loading at least 100 ng RNA per sample and using the qScript cDNA Supermix (Quantabio). Briefly, in a final volume of 20 μL, the qScript Supermix (5X) was mixed with RNA template and molecular biology grade water (Corning, 46-000-CV). cDNA was synthesized using the following PCR program: 5 minutes at 25°C, 30 minutes at 42°C, then 5 minutes at 85°C. For all quantitative real-time PCR (qPCR) reactions, cDNA was diluted to a final concentration of 5 ng/μL.

#### Quantitative real-time PCR (qPCR)

qPCR was performed by adding equal amounts of cDNA template to a mix of validated gene-specific primers, molecular biology grade water, and PerfeCta SYBR Green FastMix Low ROX (2X) (Quantabio, 84073). qPCR was performed with 3 technical replicates using a QuantStudio3 Real-Time PCR system (Thermo Fisher). Relative expression was calculated as fold change using the 2^-Δ ΔCt^ method. Primer sequences can be found in Tables 1 and 2.

**Table 1.**
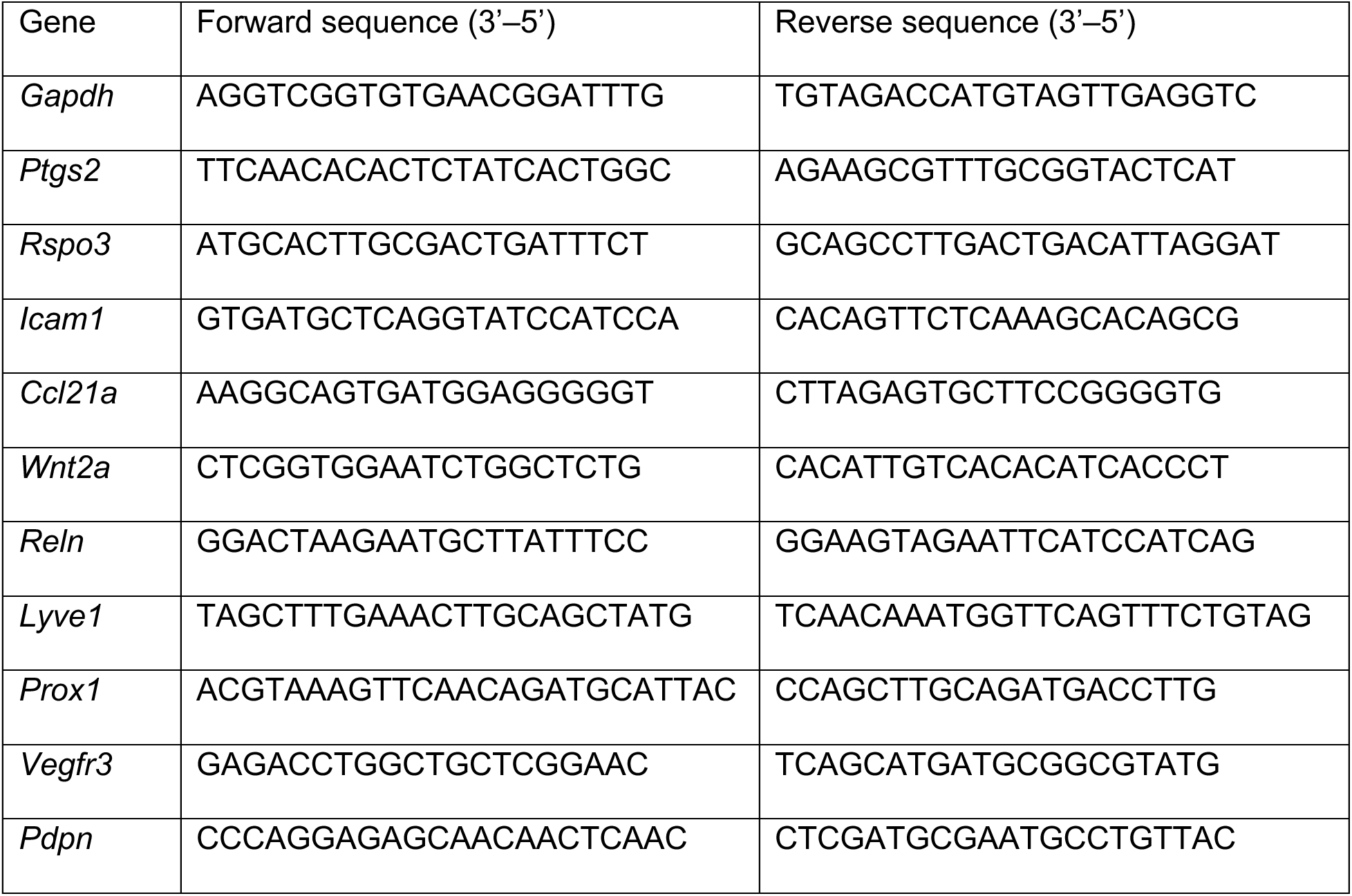
RT-qPCR primer sequences for mouse genes

**Table 2.**
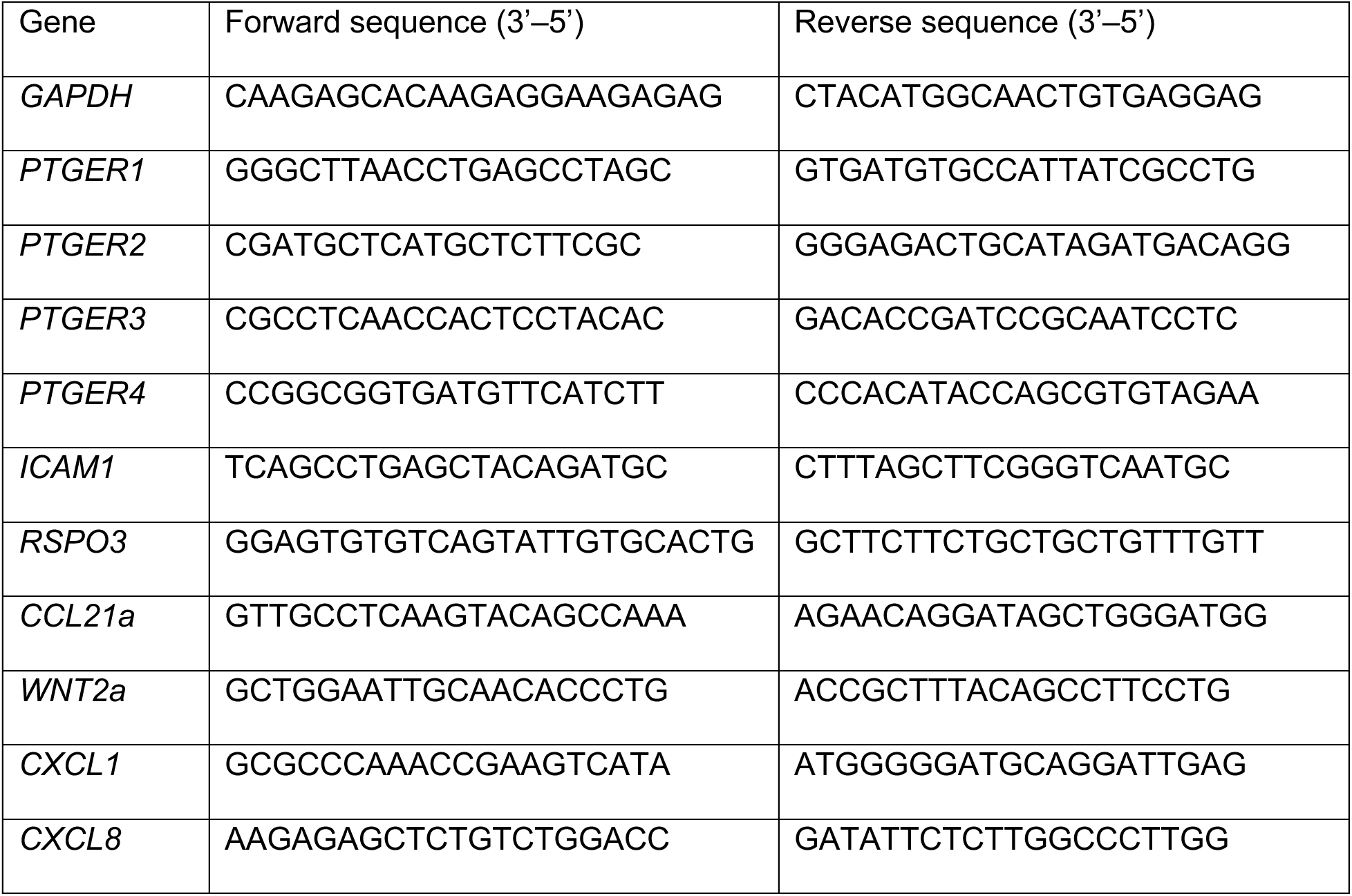
RT-qPCR primer sequences for human genes

#### RT^2^ Profiler PCR assay

cDNA from isolated Hdc^GFP^ cells was used in the RT² Profiler™ PCR Array Mouse Chemokines & Receptors cDNA (Qiagen). Manufacturer protocols were followed.

#### Bulk RNA sequencing and data processing

Isolated total RNA from HDLECs was measured for purity and integrity (RIN ≥ 8) on a 2100 Bioanalyzer (Agilent) then sent to the JP Sulzberger Columbia Genome Center for sequencing. Poly-A pulldown was used to enrich mRNAs from samples, then libraries were constructed using Illumina TruSeq chemistry. RNA-seq experiments were run on an Illumina NovaSeq 6000 with a sequencing depth of 20 million reads per sample and targeted number of paired-end 100bp reads for each sample. RTA (Illumina) was used for base calling and bcl2fastq2 version 2.19 (Illumina) was used to create FASTQ files and trim adaptors. Pseudoalignment was performed to a kallisto index created from transcriptomes (Ensembl v96, GRGm38.p6) using kallisto version 0.44.0 (Bray et al., 2016). Differentially expressed genes were determined using DESeq2 version 1.24.0 (Love et al., 2014). Gene set enrichment analysis was performed using clusterProfiler version 4.0 (Wu et al., 2021).

#### Statistical Analysis

Statistical testing was performed using GraphPad Prism 7 software (GraphPad Software Inc.). For experiments with two groups, unless otherwise specified, the differences between the means were compared using the two-sided Student’s *t*-test. For experiments with three or more groups, one-way ANOVA with post hoc Tukey’s multiple comparisons were performed. For survival experiments, the Kaplan-Meier simple survival analysis was used. Significance was defined as a *p* value < 0.05.

## DATA AND SOFTWARE AVAILABILITY

The RNA-seq dataset generated during this study is available at Gene Expression Omnibus (GEO: GSE223854).

This paper does not report original code.

Any additional information required to reanalyze the data reported in this paper is available from the lead contact upon request.

## REFERENCES

1. Abud, H.E., Watson, N., and Heath, J.K. (2005). Growth of intestinal epithelium in organ culture is dependent on EGF signalling. Exp Cell Res 303, 252–262.

2. Antanaviciute, A., Kusumbe, A., and Simmons, A. (2022). Lymphatic endothelia stakeout cryptic stem cells. Cell Stem Cell 29, 1292–1293.

3. Augustin, H.G., and Koh, G.Y. (2017). Organotypic vasculature: From descriptive heterogeneity to functional pathophysiology. Science 357.

4. Azam, S.H., Smith, M., Somasundaram, V., and Pecot, C.V. (2018). Incorporating Pericytes into an Endothelial Cell Bead Sprouting Assay. J Vis Exp.

5. Bain, C.C., and Mowat, A.M. (2014). Macrophages in intestinal homeostasis and inflammation. Immunol Rev 260, 102–117.

6. Bankhead, P., Loughrey, M.B., Fernandez, J.A., Dombrowski, Y., McArt, D.G., Dunne, P.D., McQuaid, S., Gray, R.T., Murray, L.J., Coleman, H.G., et al. (2017). QuPath: Open source software for digital pathology image analysis. Sci Rep 7, 16878.

7. Biswas, L., Chen, J., De Angelis, J., Singh, A., Owen-Woods, C., Ding, Z., Pujol, J.M., Kumar, N., Zeng, F., Ramasamy, S.K., et al. (2023). Lymphatic vessels in bone support regeneration after injury. Cell 186, 382–397 e324.

8. Blanpain, C., and Fuchs, E. (2014). Stem cell plasticity. Plasticity of epithelial stem cells in tissue regeneration. Science 344, 1242281.

9. Booth, C., Tudor, G., Tudor, J., Katz, B.P., and MacVittie, T.J. (2012). Acute gastrointestinal syndrome in high-dose irradiated mice. Health Phys 103, 383–399.

10. Bray, N.L., Pimentel, H., Melsted, P., and Pachter, L. (2016). Near-optimal probabilistic RNA-seq quantification. Nat Biotechnol 34, 525–527.

11. Chen, X., Deng, H., Churchill, M.J., Luchsinger, L.L., Du, X., Chu, T.H., Friedman, R.A., Middelhoff, M., Ding, H., Tailor, Y.H., et al. (2017). Bone Marrow Myeloid Cells Regulate Myeloid-Biased Hematopoietic Stem Cells via a Histamine-Dependent Feedback Loop. Cell Stem Cell 21, 747–760 e747.

12. Choe, K., Jang, J.Y., Park, I., Kim, Y., Ahn, S., Park, D.Y., Hong, Y.K., Alitalo, K., Koh, G.Y., and Kim, P. (2015). Intravital imaging of intestinal lacteals unveils lipid drainage through contractility. J Clin Invest 125, 4042–4052.

13. Couter, C.J., and Surana, N.K. (2016). Isolation and Flow Cytometric Characterization of Murine Small Intestinal Lymphocytes. J Vis Exp.

14. Crittenden, S., Goepp, M., Pollock, J., Robb, C.T., Smyth, D.J., Zhou, Y., Andrews, R., Tyrrell, V., Gkikas, K., Adima, A., et al. (2021). Prostaglandin E2 promotes intestinal inflammation via inhibiting microbiota-dependent regulatory T cells. Sci Adv 7.

15. Cursiefen, C., Chen, L., Borges, L.P., Jackson, D., Cao, J., Radziejewski, C., D’Amore, P.A., Dana, M.R., Wiegand, S.J., and Streilein, J.W. (2004). VEGF-A stimulates lymphangiogenesis and hemangiogenesis in inflammatory neovascularization via macrophage recruitment. J Clin Invest 113, 1040–1050.

16. De Schepper, S., Verheijden, S., Aguilera-Lizarraga, J., Viola, M.F., Boesmans, W., Stakenborg, N., Voytyuk, I., Schmidt, I., Boeckx, B., Dierckx de Casterle, I., et al. (2018). Self-Maintaining Gut Macrophages Are Essential for Intestinal Homeostasis. Cell 175, 400–415 e413.

17. Fu, N., Wu, F., Jiang, Z., Kim, W., Ruan, T., Malagola, E., Ochiai, Y., Napoles, O.C., Valenti, G., White, R.A., et al. (2021). Acute Intestinal Inflammation Depletes/Recruits Histamine-Expressing Myeloid Cells From the Bone Marrow Leading to Exhaustion of MB-HSCs. Cell Mol Gastroenterol Hepatol 11, 1119–1138.

18. Goto, N., Goto, S., Imada, S., Hosseini, S., Deshpande, V., and Yilmaz, O.H. (2022). Lymphatics and fibroblasts support intestinal stem cells in homeostasis and injury. Cell Stem Cell 29, 1246–1261 e1246.

19. Green, D.E., and Rubin, C.T. (2014). Consequences of irradiation on bone and marrow phenotypes, and its relation to disruption of hematopoietic precursors. Bone 63, 87–94.

20. Guo, S., Nighot, M., Al-Sadi, R., Alhmoud, T., Nighot, P., and Ma, T.Y. (2015). Lipopolysaccharide Regulation of Intestinal Tight Junction Permeability Is Mediated by TLR4 Signal Transduction Pathway Activation of FAK and MyD88. J Immunol 195, 4999-5010.

21. Harnack, C., Berger, H., Antanaviciute, A., Vidal, R., Sauer, S., Simmons, A., Meyer, T.F., and Sigal, M. (2019). R-spondin 3 promotes stem cell recovery and epithelial regeneration in the colon. Nat Commun 10, 4368.

22. Hull, M.A., Ko, S.C., and Hawcroft, G. (2004). Prostaglandin EP receptors: targets for treatment and prevention of colorectal cancer? Mol Cancer Ther 3, 1031–1039.

23. Ishikawa, T.O., and Herschman, H.R. (2006). Conditional knockout mouse for tissue-specific disruption of the cyclooxygenase-2 (Cox-2) gene. Genesis 44, 143–149.

24. Ishikawa, T.O., Oshima, M., and Herschman, H.R. (2011). Cox-2 deletion in myeloid and endothelial cells, but not in epithelial cells, exacerbates murine colitis. Carcinogenesis 32, 417–426.

25. Jarde, T., Chan, W.H., Rossello, F.J., Kaur Kahlon, T., Theocharous, M., Kurian Arackal, T., Flores, T., Giraud, M., Richards, E., Chan, E., et al. (2020). Mesenchymal Niche-Derived Neuregulin-1 Drives Intestinal Stem Cell Proliferation and Regeneration of Damaged Epithelium. Cell Stem Cell 27, 646–662 e647.

26. Katoh, H., Wang, D., Daikoku, T., Sun, H., Dey, S.K., and Dubois, R.N. (2013). CXCR2-expressing myeloid-derived suppressor cells are essential to promote colitis-associated tumorigenesis. Cancer Cell 24, 631–644.

27. Kim, M., Galan, C., Hill, A.A., Wu, W.J., Fehlner-Peach, H., Song, H.W., Schady, D., Bettini, M.L., Simpson, K.W., Longman, R.S., et al. (2018). Critical Role for the Microbiota in CX3CR1(+) Intestinal Mononuclear Phagocyte Regulation of Intestinal T Cell Responses. Immunity 49, 151–163 e155.

28. Liang, L., Shen, L., Fu, G., Yao, Y., Li, G., Deng, Y., Zhang, H., Zhou, M., Yang, W., Hua, G., et al. (2020). Regulation of the regeneration of intestinal stem cells after irradiation. Ann Transl Med 8, 1063.

29. Love, M.I., Huber, W., and Anders, S. (2014). Moderated estimation of fold change and dispersion for RNA-seq data with DESeq2. Genome Biol 15, 550.

30. Machnik, A., Neuhofer, W., Jantsch, J., Dahlmann, A., Tammela, T., Machura, K., Park, J.K., Beck, F.X., Muller, D.N., Derer, W., et al. (2009). Macrophages regulate salt-dependent volume and blood pressure by a vascular endothelial growth factor-C-dependent buffering mechanism. Nat Med 15, 545–552.

31. McCarthy, N., Kraiczy, J., and Shivdasani, R.A. (2020a). Cellular and molecular architecture of the intestinal stem cell niche. Nat Cell Biol 22, 1033–1041.

32. McCarthy, N., Manieri, E., Storm, E.E., Saadatpour, A., Luoma, A.M., Kapoor, V.N., Madha, S., Gaynor, L.T., Cox, C., Keerthivasan, S., et al. (2020b). Distinct Mesenchymal Cell Populations Generate the Essential Intestinal BMP Signaling Gradient. Cell Stem Cell 26, 391–402 e395.

33. Miyoshi, H., VanDussen, K.L., Malvin, N.P., Ryu, S.H., Wang, Y., Sonnek, N.M., Lai, C.W., and Stappenbeck, T.S. (2017). Prostaglandin E2 promotes intestinal repair through an adaptive cellular response of the epithelium. EMBO J 36, 5–24.

34. Muley, A., Odaka, Y., Lewkowich, I.P., Vemaraju, S., Yamaguchi, T.P., Shawber, C., Dickie, B.H., and Lang, R.A. (2017). Myeloid Wnt ligands are required for normal development of dermal lymphatic vasculature. PLoS One 12, e0181549.

35. Murtomaki, A., Uh, M.K., Choi, Y.K., Kitajewski, C., Borisenko, V., Kitajewski, J., and Shawber, C.J. (2013). Notch1 functions as a negative regulator of lymphatic endothelial cell differentiation in the venous endothelium. Development 140, 2365–2376.

36. Nandi, P., Girish, G.V., Majumder, M., Xin, X., Tutunea-Fatan, E., and Lala, P.K. (2017). PGE2 promotes breast cancer-associated lymphangiogenesis by activation of EP4 receptor on lymphatic endothelial cells. BMC Cancer 17, 11.

37. Niec, R.E., Chu, T., Schernthanner, M., Gur-Cohen, S., Hidalgo, L., Pasolli, H.A., Luckett, K.A., Wang, Z., Bhalla, S.R., Cambuli, F., et al. (2022). Lymphatics act as a signaling hub to regulate intestinal stem cell activity. Cell Stem Cell 29, 1067–1082 e1018.

38. Nighot, M., Al-Sadi, R., Guo, S., Rawat, M., Nighot, P., Watterson, M.D., and Ma, T.Y. (2017). Lipopolysaccharide-Induced Increase in Intestinal Epithelial Tight Permeability Is Mediated by Toll-Like Receptor 4/Myeloid Differentiation Primary Response 88 (MyD88) Activation of Myosin Light Chain Kinase Expression. Am J Pathol 187, 2698–2710.

39. Ocansey, D.K.W., Pei, B., Xu, X., Zhang, L., Olovo, C.V., and Mao, F. (2021). Cellular and molecular mediators of lymphangiogenesis in inflammatory bowel disease. J Transl Med 19, 254.

40. Ogasawara, R., Hashimoto, D., Kimura, S., Hayase, E., Ara, T., Takahashi, S., Ohigashi, H., Yoshioka, K., Tateno, T., Yokoyama, E., et al. (2018). Intestinal Lymphatic Endothelial Cells Produce R-Spondin3. Sci Rep 8, 10719.

41. Palikuqi, B., Nguyen, D.T., Li, G., Schreiner, R., Pellegata, A.F., Liu, Y., Redmond, D., Geng, F., Lin, Y., Gomez-Salinero, J.M., et al. (2020). Adaptable haemodynamic endothelial cells for organogenesis and tumorigenesis. Nature 585, 426–432.

42. Palikuqi, B., Rispal, J., Reyes, E.A., Vaka, D., Boffelli, D., and Klein, O. (2022). Lymphangiocrine signals are required for proper intestinal repair after cytotoxic injury. Cell Stem Cell 29, 1262–1272 e1265.

43. Rafii, S., Butler, J.M., and Ding, B.S. (2016). Angiocrine functions of organ-specific endothelial cells. Nature 529, 316–325.

44. Roulis, M., Kaklamanos, A., Schernthanner, M., Bielecki, P., Zhao, J., Kaffe, E., Frommelt, L.S., Qu, R., Knapp, M.S., Henriques, A., et al. (2020). Paracrine orchestration of intestinal tumorigenesis by a mesenchymal niche. Nature 580, 524–529.

45. Saha, S., Aranda, E., Hayakawa, Y., Bhanja, P., Atay, S., Brodin, N.P., Li, J., Asfaha, S., Liu, L., Tailor, Y., et al. (2016). Macrophage-derived extracellular vesicle-packaged WNTs rescue intestinal stem cells and enhance survival after radiation injury. Nat Commun 7, 13096.

46. Sato, T., Vries, R.G., Snippert, H.J., van de Wetering, M., Barker, N., Stange, D.E., van Es, J.H., Abo, A., Kujala, P., Peters, P.J., et al. (2009). Single Lgr5 stem cells build crypt-villus structures in vitro without a mesenchymal niche. Nature 459, 262–265.

47. Schindelin, J., Arganda-Carreras, I., Frise, E., Kaynig, V., Longair, M., Pietzsch, T., Preibisch, S., Rueden, C., Saalfeld, S., Schmid, B., et al. (2012). Fiji: an open-source platform for biological-image analysis. Nat Methods 9, 676–682.

48. Shaw, T.N., Houston, S.A., Wemyss, K., Bridgeman, H.M., Barbera, T.A., Zangerle-Murray, T., Strangward, P., Ridley, A.J.L., Wang, P., Tamoutounour, S., et al. (2018). Tissue-resident macrophages in the intestine are long lived and defined by Tim-4 and CD4 expression. J Exp Med 215, 1507–1518.

49. Smith, P.D., Smythies, L.E., Shen, R., Greenwell-Wild, T., Gliozzi, M., and Wahl, S.M. (2011). Intestinal macrophages and response to microbial encroachment. Mucosal Immunol 4, 31–42.

50. Tacconi, C., Correale, C., Gandelli, A., Spinelli, A., Dejana, E., D’Alessio, S., and Danese, S. (2015). Vascular endothelial growth factor C disrupts the endothelial lymphatic barrier to promote colorectal cancer invasion. Gastroenterology 148, 1438–1451 e1438.

51. Tessner, T.G., Muhale, F., Riehl, T.E., Anant, S., and Stenson, W.F. (2004). Prostaglandin E2 reduces radiation-induced epithelial apoptosis through a mechanism involving AKT activation and bax translocation. J Clin Invest 114, 1676–1685.

52. Walden, T.L., Jr., Patchen, M., and Snyder, S.L. (1987). 16,16-Dimethyl prostaglandin E2 increases survival in mice following irradiation. Radiat Res 109, 440–448.

53. Wu, T., Hu, E., Xu, S., Chen, M., Guo, P., Dai, Z., Feng, T., Zhou, L., Tang, W., Zhan, L., et al. (2021). clusterProfiler 4.0: A universal enrichment tool for interpreting omics data. Innovation (Camb) 2, 100141.

54. Xu, J., Zhang, C., He, Y., Wu, H., Wang, Z., Song, W., Li, W., He, W., Cai, S., and Zhan, W. (2012). Lymphatic endothelial cell-secreted CXCL1 stimulates lymphangiogenesis and metastasis of gastric cancer. Int J Cancer 130, 787–797.

55. Yan, K.S., Janda, C.Y., Chang, J., Zheng, G.X.Y., Larkin, K.A., Luca, V.C., Chia, L.A., Mah, A.T., Han, A., Terry, J.M., et al. (2017). Non-equivalence of Wnt and R-spondin ligands during Lgr5(+) intestinal stem-cell self-renewal. Nature 545, 238–242.

56. Yang, C., Yu, H., Chen, R., Tao, K., Jian, L., Peng, M., Li, X., Liu, M., and Liu, S. (2019). CXCL1 stimulates migration and invasion in ER-negative breast cancer cells via activation of the ERK/MMP2/9 signaling axis. Int J Oncol 55, 684–696.

57. Yang, X.D., Ai, W., Asfaha, S., Bhagat, G., Friedman, R.A., Jin, G., Park, H., Shykind, B., Diacovo, T.G., Falus, A., et al. (2011). Histamine deficiency promotes inflammation-associated carcinogenesis through reduced myeloid maturation and accumulation of CD11b+Ly6G+ immature myeloid cells. Nat Med 17, 87–95.

58. Zecharia, A.Y., Yu, X., Gotz, T., Ye, Z., Carr, D.R., Wulff, P., Bettler, B., Vyssotski, A.L., Brickley, S.G., Franks, N.P., et al. (2012). GABAergic inhibition of histaminergic neurons regulates active waking but not the sleep-wake switch or propofol-induced loss of consciousness. J Neurosci 32, 13062–13075.

