## Supplemental figure legends for "Immature myeloid cells are indispensable for intestinal regeneration post irradiation injury"

**SUPPLEMENTAL FIGURES**

**Supplementary Figure S1: Granulocytic Hdc+ IMCs invade the irradiated intestine.**

**Supplementary Figure S2: PGE2 promotes intestinal epithelial proliferation.**

**Supplementary Figure S3: PGE2 promotes lymphatic sprouting and expression of regenerative factors.**

**Supplementary Figure S4: Microbial signals promote LEC recruitment of Hdc+ IMCs.**

### SUPPLEMENTARY FIGURE LEGENDS

**Supplementary Figure 1: Granulocytic Hdc+ IMCs invade the irradiated intestine.**

**A.** Representative 100x images of $Hdc^{GFP}$ visualization from the proximal jejunum of non-irradiated $Hdc^{GFP}$ mice or mice after 12 Gy WB-IR. **B.** Representative FACS plots of live ($DAPI^-$) intestinal cells from non-irradiated $Hdc^{GFP}$ mice or mice 3 and 10 days after 12 Gy WB-IR. **C.** Bar plot showing quantification of $Hdc^{GFP+}$ cells of images in (**A**). (n=2 or 3), HPF: high powered field. **D.** Bar plot showing quantification of (**B**). (n=3 or 4). **E.** Representative FACs plot of $DAPI^-$, $CD45^+$, $Hdc^{GFP+}$ cells isolated from the intestine of $Hdc^{GFP}$ mice after 12 Gy WB-IR. **F.** Representative FACs plot of $DAPI^-$, $CD45^+$, cells isolated from the small intestine of $Hdc^{CreERT2}$ mice 3 days after 12 Gy WB-IR. **G and H)** Bar plots showing quantifications of (**E**). (n=3 or 4).

Scale bar = 50 um.

Bar graph data are mean ± SEM.

Statistical analysis was performed using an Ordinary one-way ANOVA with multiple comparisons to each group (**C, D**) or a two-sided Student’s t test (**G, H**).

**Supplementary Figure 2: PGE2 promotes intestinal epithelial proliferation.**

**A.** Representative 200x images of intestinal organoids in culture treated with various concentrations of dmPGE2 with or without EP4i. **B**. Bar plot showing quantification of the average sphere number of (**A**). (n=3). **C.** Bar plot showing quantification of the average sphere size of (**A**). (n=3). **D.** Scheme of dmPGE2 treatment experiment. I.P. = intraperitoneal injection of vehicle or dmPGE2. **E.** Representative 100x images of H&E staining from the proximal jejunum of $Hdc^{GFP}$ mice treated with vehicle or dmPGE2 for 3 days. **F.** Bar plot showing quantification of the average villus length of images in (**D**). (n=3). **G.** Representative images of BrdU staining from the proximal jejunum of $Hdc^{GFP}$ mice treated with vehicle or dmPGE2. **H.** Bar plot showing quantification of BrdU staining from (**G**). (n=3).

Scale bar = 50 um.

Bar graph data are mean ± SEM.

Statistical analysis was performed using an Ordinary one-way ANOVA with multiple comparisons to Ctrl (**B, C**) or a two-sided Student's t test (**E, G**).

**Supplementary Figure 3: PGE2 promotes lymphatic sprouting and expression of regenerative factors.**

**A.** qPCR analysis of sorted live $CD45^-$ $EpCAM^-$ $CD31^+$ $CD90.2^{+/-}$ cells for LEC-related genes. (n=4). **B.** Bar plot showing quantification of the percentage of $CD90^+$ LECs ($CD31^+$ $CD90.2^+$) out of all $DAPI^-$ $CD45^-$ $EpCAM^-$ cells from the small intestines of $Hdc^{CreERT2}$; $R26^{DTA}$ mice after 12 Gy WB-IR, that received adoptive transfer of $Hdc^{GFP-}$ bone marrow cells ($GFP^-$ A.T.) or $Hdc^{GFP+}$ bone marrow cells ($GFP^+$ A.T.) from healthy $Hdc^{GFP}$ mice twice after irradiation. See figure (**1I**) for scheme. (n=3). **C.** qPCR analysis of *Ptger* genes of sorted LECs from the intestines of mice. (n=3). **D.** Pathway analysis of differentially expressed genes obtained from bulk RNA sequencing of cultured HDLECs treated with PGE2 compared to control. (n=4). **E**. Representative 200x images of HDLEC cells on microcarrier beads treated with vehicle, dmPGE2, or dmPGE2 with EP4i. **F.** Bar plot showing quantification of the average sprout length of (**E**). (n=3). **G.** qPCR analysis of activation and regenerative genes of cultured HDLECs treated with vehicle, dmPGE2, or dmPGE2 with EP4i for 6 hours. (n=3). **H.** ELISA analysis of supernatant RSPO3 isolated from cultured HDLECs treated with vehicle, dmPGE2, or dmPGE2 + EP4i for 16 hours. (n=3). **I**. qPCR analysis of activation and regenerative genes of healthy mice treated with vehicle or dmPGE2 for 3 days. (n=3).

Scale bars = 50 um.

Bar graph data are mean ± SEM.

Statistical analysis was performed using an Ordinary one-way ANOVA with multiple comparisons to each group (**E, G**) or a two-sided Student's t test (**A, F, H**).

**Supplementary Figure 4: Microbial signals promote LEC recruitment of Hdc+ IMCs.**

**A.** qPCR for *CXCL1* and *CXCL8* in cultured HDLEC treated with various LPS concentrations for 6 hours. (n=3). **B.** Scheme of Transwell migration assay. **D.** Quantification of results of Transwell migration assay. (n=3). CXCR2i: SB225002. MEK1/2i: U0126.

Bar graph data are mean ± SEM.

Statistical analysis was performed using an Ordinary one-way ANOVA with multiple comparisons to each group (**A, D**).
