## Supplemental figures for "Immature myeloid cells are indispensable for intestinal regeneration post irradiation injury"

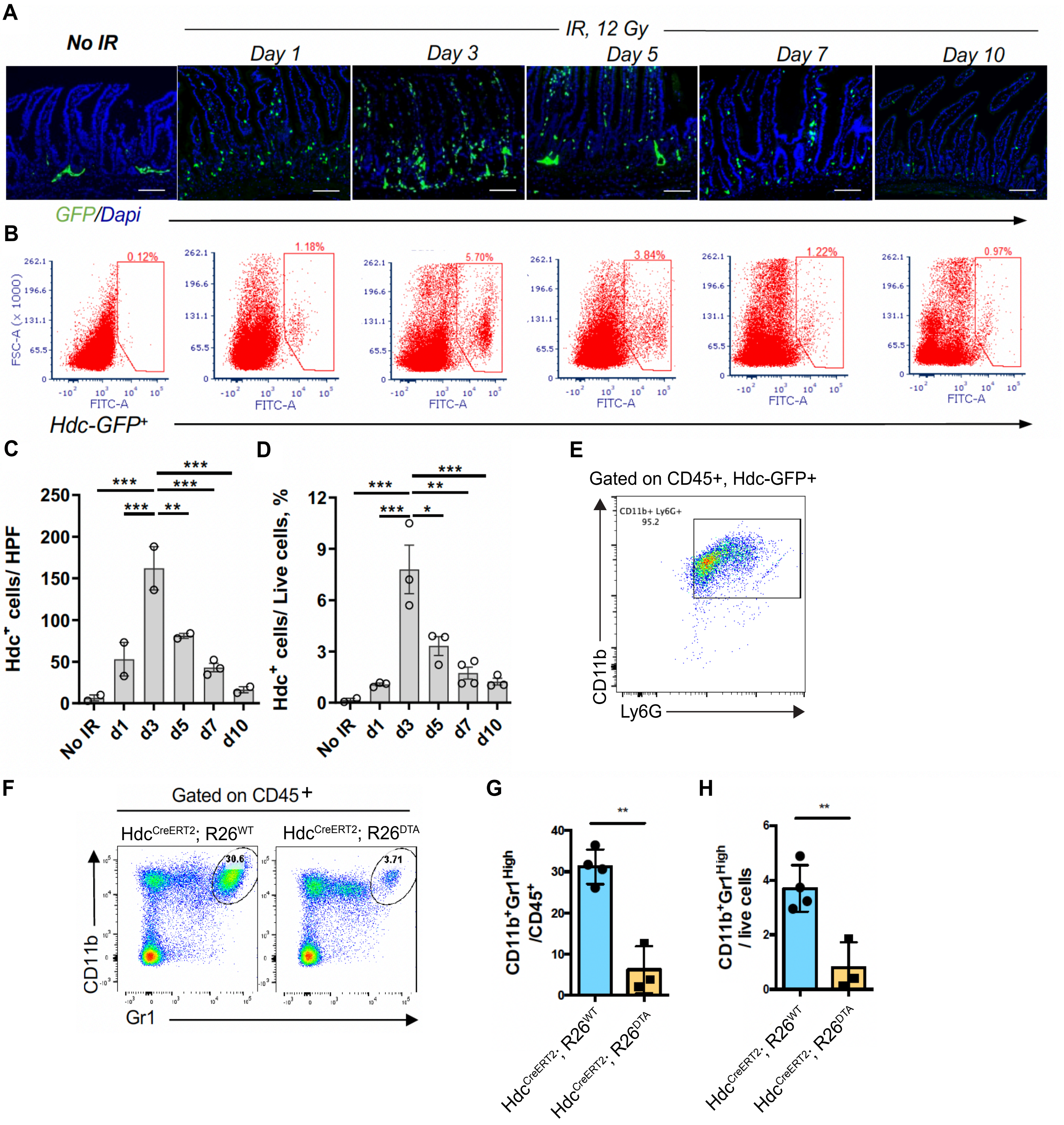


**Figure S1: Granulocytic Hdc+ IMCs invade the irradiated intestine.**

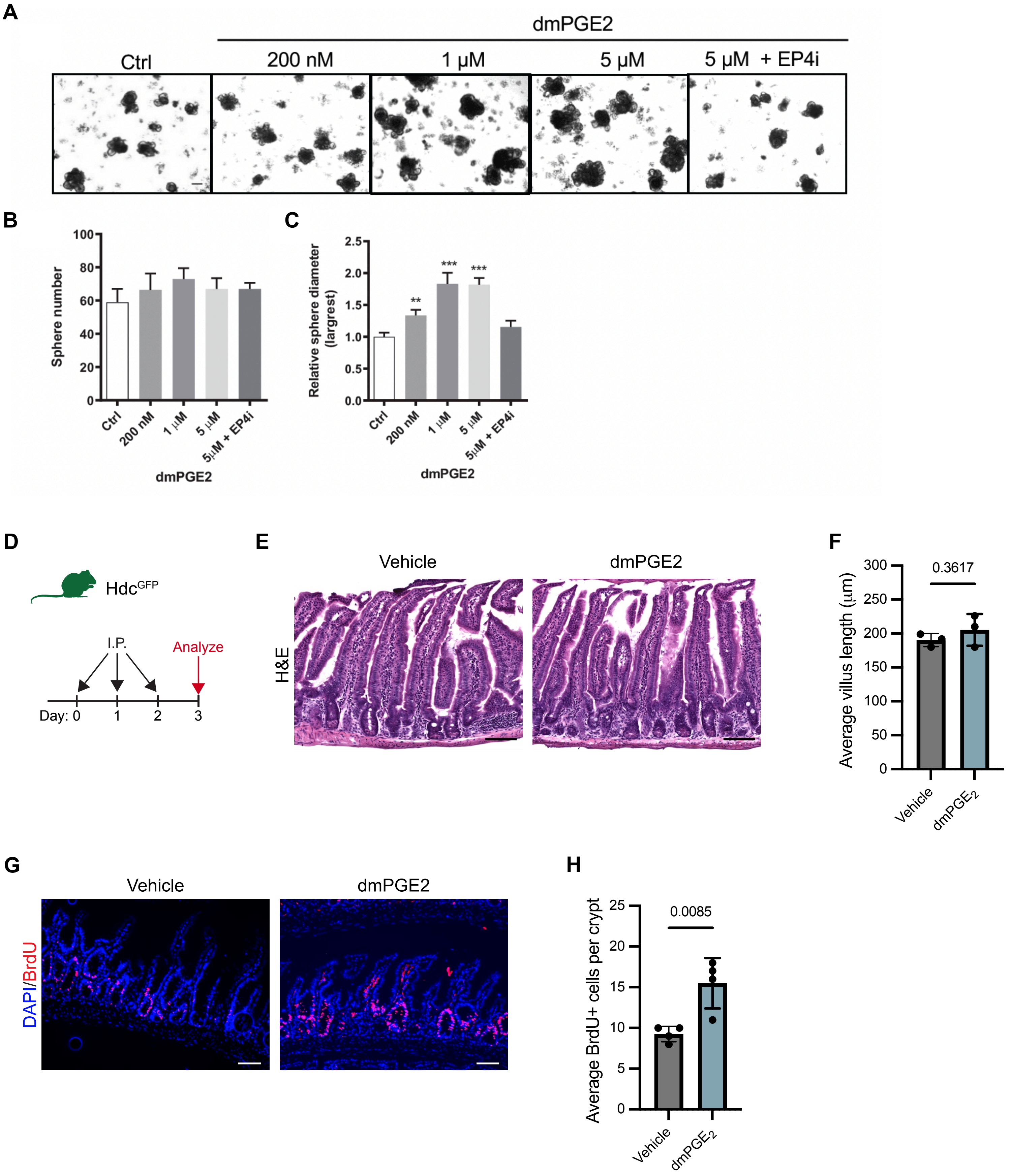


**Figure S2: PGE2 promotes intestinal epithelial proliferation.**

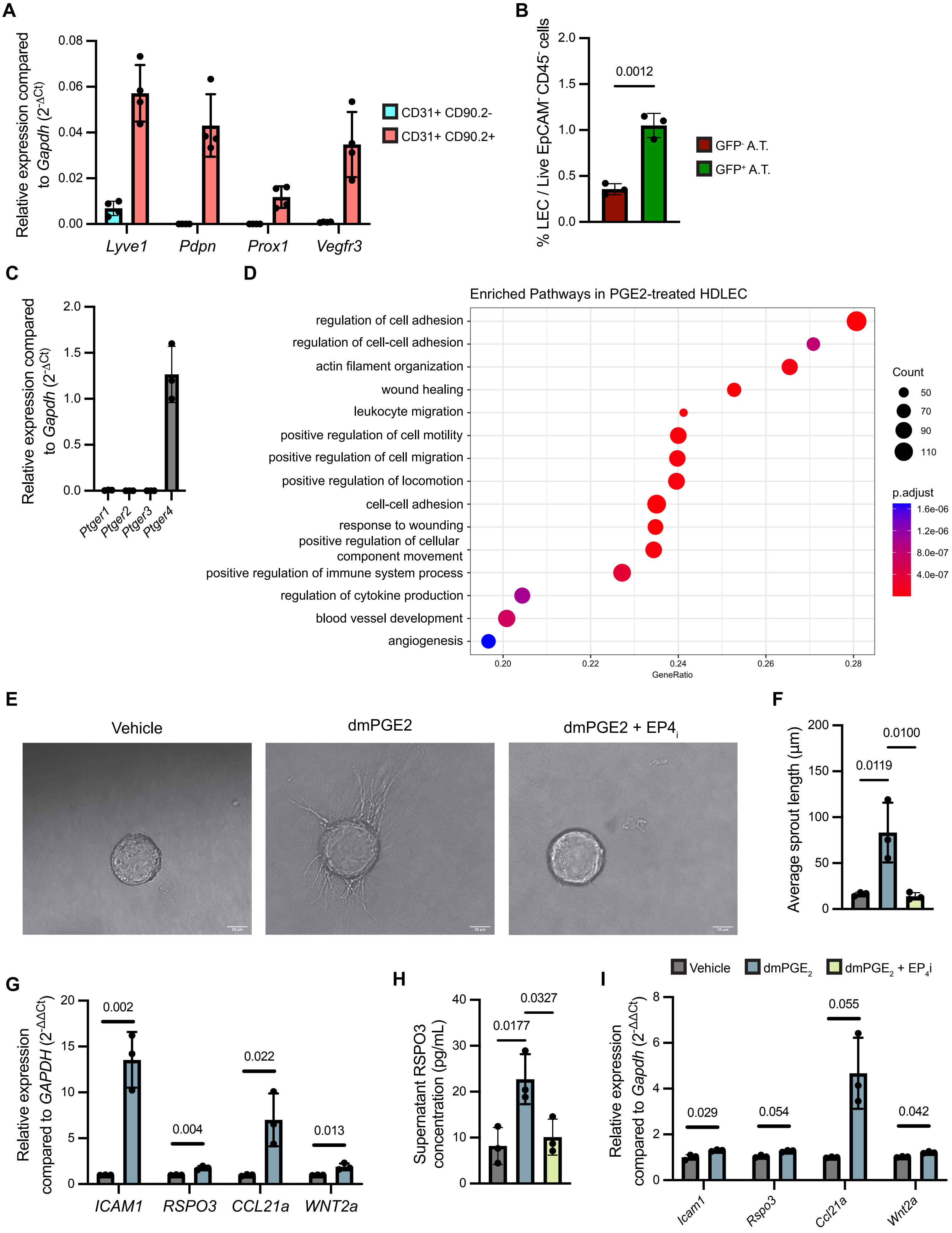


**Figure S3: PGE2 promotes lymphatic sprouting and expression of regenerative factors.**

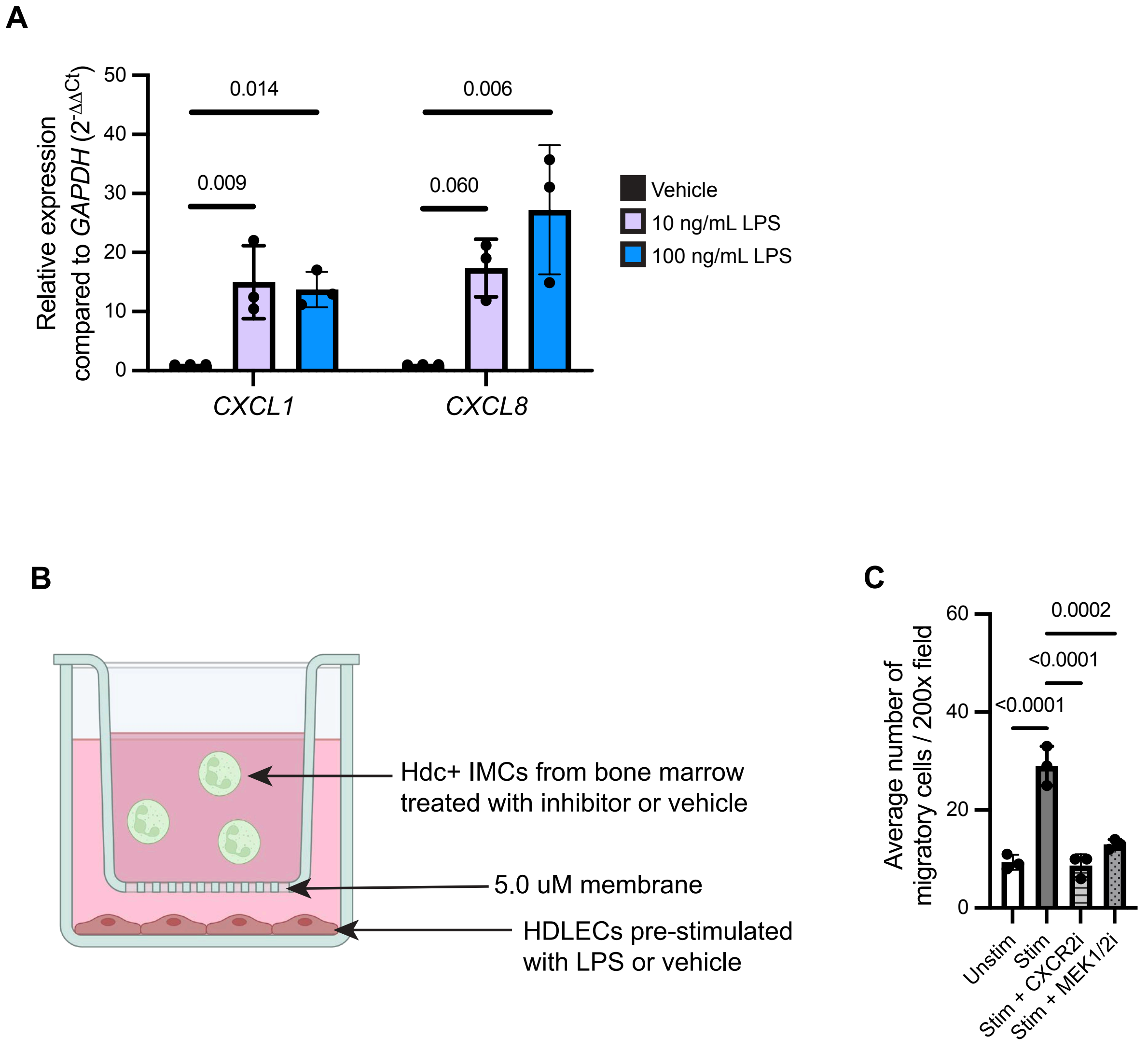


**Figure S4: Microbial signals promote LEC recruitment of Hdc+ IMCs.**
